# Syngap1 Regulates Cortical Circuit Assembly by Controlling Membrane Excitability

**DOI:** 10.1101/2022.12.06.519295

**Authors:** Vineet Arora, Sheldon Michaelson, Massimiliano Aceti, Murat Kilinic, Courtney Miller, Gavin Rumbaugh

**Affiliations:** Departments of Neuroscience and Molecular Medicine, UF Scripps Biomedical Research Institute (Formerly Scripps Florida), Jupiter, FL. 33458; Skaggs Graduate School of Chemical and Biological Sciences, Scripps Research, Jupiter, FL 33458; Oncology R&D, AstraZeneca, Cambridge, Cambridgeshire CB2 0AA, UK; Alkahest, Inc., 125 Shoreway Road, Suite D, San Carlos, CA 94070

**Keywords:** gene, development, activity, circuit, autism, cortex, synapse, Syngap1

## Abstract

Gene expression intersects with neural activity to produce cortical circuits during brain development. However, the cell biological mechanisms linking gene expression to activity-dependent cortical circuit assembly remain unclear. Here, we demonstrate in mice that a newly discovered function of the neurodevelopmental disorder gene, *Syngap1*, is to cell-autonomously control intrinsic membrane excitability (IME) in developing cortical glutamatergic neurons. *Syngap1* regulation of IME was mechanistically linked to wiring of a cortical circuit motif required for sensory processing and behavioral action. Restoring depressed IME in *Syngap1* deficient neurons through genetic targeting of hyper-functional potassium currents unleashed deficient dendritic morphogenesis in upper lamina sensory cortex pyramidal neurons. Furthermore, enhancing dendritic morphogenesis was sufficient to stimulate assembly of translaminar feed-forward excitatory circuit motifs. Thus, *Syngap1* promotes excitatory circuit assembly during cortical development by maintaining IME in a range that enables trophic neuronal activity to maximize pyramidal cell somatodendritic maturation and subsequent synapse formation.

**Highlights:**

- *Syngap1* cell-autonomously tunes cortical pyramidal neuron IME *in vivo*
- *Syngap1*-IME is regulated in part by control of neuronal potassium currents
- *Syngap1* enhancement of IME drives dendritic maturation in pyramidal cells
- *Syngap1* tuning of IME-regulated dendritic maturation promotes circuit assembly

## Introduction

Neural dynamics within cortical circuits lead to cognitive transformations that enable adaptive behaviors (Buschman and Kastner, 2015; Constantinople and Bruno, 2011; Gallero-Salas et al., 2021; Gilad and Helmchen, 2020; Helmchen et al., 2018; MacDowell and Buschman, 2020; Robertson and Baron-Cohen, 2017; Staiger and Petersen, 2021). Genes contribute to the formation of cortical circuits through activity-dependent processes (Ebert and Greenberg, 2013). Early in development, activity is generated intrinsically within nascent circuits through genetic programs, which is a process that helps to segregate neural connectivity into organized pathways, or circuits (Ackman et al., 2012; Luhmann and Khazipov, 2018). In later stages of development and throughout adulthood, the source of activity within nascent circuits is shifted toward sensory experience (Ribic, 2020). Activity through sensory experience acts to refine the coarse patterns of neural connectivity initially built through intrinsic mechanisms. Recent technical advances have led to the discovery of a previously unappreciated number of neurotransmitter-specific neuronal subtypes even within the same brain region (Tasic et al., 2018; Yao et al., 2021). By providing a basic “parts list”, this advance has led to insights into how complex and highly specified cortical circuits form during development. However, with this new level of circuit element granularity, it is important to study molecular and cellular mechanisms of cortical development within functionally defined circuit motifs known to regulate specific aspects of cognitive processing and behavioral control.

Cortical sensory processing is required for perceptual decision making that underlies goal-directed behavioral action (Staiger and Petersen, 2021). A function of primary sensory cortex circuitry is to integrate ascending sensory information arriving from the periphery with top-down signals for current internal state and prior relevant experience (Feldmeyer et al., 2013). For this integration to occur, sensory information originating in sub-cortical areas must be routed into and throughout the neocortex, which occurs through a series of discrete synaptic circuit motifs that function locally within distributed sensorimotor systems. Principle among these circuits are the feed-forward motifs in upper lamina primary sensory areas. Feed-forward excitatory motifs are defined by a high density of layer 4 (L4) stellate neurons that receive sensory thalamus input, which then project directly to L2/3 pyramidal neurons within the same cortical column (Petersen and Crochet, 2013). A crucial neurophysiological function that these circuits subserve is to promote cortical representations of peripherally generated sensory signals (Romo and Rossi-Pool, 2020). Activity within these representations is important for behavioral control. For example, the activity within somatosensory cortex (SSC) neurons can predict the intensity of peripheral tactile stimuli (de Lafuente and Romo, 2005, 2006) and behavioral choice (Kwon et al., 2016; Stüttgen and Schwarz, 2008). Moreover, lowering spiking activity of pyramidal neurons in SSC is sufficient to alter sensory-informed goal-directed behavioral action in expertly trained animals (Sachidhanandam et al., 2013). Spike coding in SSC feed-forward circuits is thought to aid in decision-making by driving downstream cortical regions involved in choice-related motor responses (Kwon *et al*., 2016; Romo and Rossi-Pool, 2020). As such, identifying the molecular and cellular mechanisms that shape the assembly of these feedforward sensory cortex circuit motifs may reveal neurodevelopmental substrates of cognitive maturation.

Within the cortex, activity dependent processes shape the formation of circuit motifs important for cognitive processing and behavioral control. What are the genetic mechanisms that promote the intrinsic assembly of cortical circuits and how do these genes gate neural activity that instruct the assembly process? It has been argued that highly penetrant genes that cause rare forms of neurodevelopmental disorders (NDDs) can promote discovery of activity dependent mechanisms that promote precise circuit assembly programs during brain development (Ebert and Greenberg, 2013). Indeed, *de novo* genetic loss-of-function single nucleotide variants (SNVs) within individual genes can lead to disorders that are defined by impaired cognitive functioning and altered sensory processing (Deciphering Developmental Disorders, 2015, 2017; Satterstrom et al., 2020). Cognitive impairment associated with these disorders, including autism spectrum disorder and schizophrenia, is thought to arise from an abnormally developing cortex that results in altered structural and functional connectivity of sensory processing circuits (Holiga et al., 2019; Hong et al., 2019; Jack, 2018; Muhle et al., 2018; Nunes et al., 2018; Tebartz van Elst et al., 2016; Yamasaki et al., 2017). Thus, in-depth study of the natural functions of genes derived from extensive NDD diagnostic sequencing campaigns can potentially reveal the neurobiological mechanisms that link activity-dependent cell-biological mechanisms to the assembly of sensory processing circuits that guide behavioral choice.

*SYNGAP1/Syngap1* (*HUMAN/Mouse*) is an attractive NDD risk gene for identifying neurodevelopmental mechanisms that may link gene expression to sensory cortex circuit assembly. Clinically, *SYNGAP1* has been repeatedly demonstrated to be a frequently mutated gene in various types of NDDs defined by impaired cognitive functioning and behavioral control (Berryer et al., 2013; Deciphering Developmental Disorders, 2015, 2017; Fu et al., 2022; Hamdan et al., 2011; Hamdan et al., 2009; Satterstrom et al., 2020). Patients with *SYNGAP1* disorders are largely defined by loss-of-function SNVs leading to genetic haploinsufficiency, which leads to reduced expression of SynGAP proteins (Holder et al., 1993; Jimenez-Gomez et al., 2019; Weldon et al., 2018). *SYNGAP1* haploinsufficiency causes intellectual impairment, epilepsy, and autistic features. The complete penetrance of pathogenic SNVs (Satterstrom *et al*., 2020), combined with the failure to identify neurotypical humans expressing pathogenic loss-of-function *SYNGAP1* SNVs from more than 100,000 samples (Karczewski et al., 2020), demonstrates that its expression during human brain development regulates cellular mechanisms that drive cognitive maturation and balanced excitability within neural systems. Importantly, *SYNGAP1* patients demonstrate clear alterations in sensory processing. They also display abnormal behaviors in response to tactile or auditory input and also exhibit alterations in neural activity in response to sensory stimulation (Cote et al., 2021a; Cote et al., 2021b; Jimenez-Gomez *et al*., 2019; Michaelson et al., 2018; Mignot et al., 2016; Parker et al., 2015; Vlaskamp et al., 2019).

Prior studies have shown that adult *Syngap1* heterozygous mice, which model genetic risk across the human patient population, have impaired tactile representations within somatosensory cortex (SSC) and tactile-guided behavioral abnormalities (Michaelson *et al*., 2018). They also exhibit impaired feed-forward excitatory circuit function in the SSC, which contributes to disrupted sensory representations within sensory processing networks. Given that sensory cortex feed-forward excitatory circuits comprise a crucial node in the larger distributed sensorimotor networks that enable perceptual decision making (Feldmeyer *et al*., 2013; Romo and Rossi-Pool, 2020; Staiger and Petersen, 2021), elucidating the biological mechanisms linking *Syngap1* expression to development of these circuit motifs may uncover neurobiological substrates of cognitive maturation. However, it remains unknown how *Syngap1* expression in these neurons regulates the assembly of these translaminar excitatory circuit motifs. Here, we identified a novel cell autonomous function of the *Syngap1* gene in developing cortex that was linked directly to the assembly of cortical feed-forward circuits: to control intrinsic membrane excitability (IME) in developing L2/3 SSC neurons. Because many highly penetrant NDD genes regulate IME in developing cortical neurons, we propose that this discovery related to *Syngap1* may be a generalizable genetic mechanism that gates activity-dependent assembly of cortical circuits and may be a point of convergence for NDD etiology.

## Results

We sought to identify the cellular mechanisms linking *Syngap1* developmental expression to assembly of SSC feed-forward excitatory circuits. L2/3 neurons from adult *Syngap1* heterozygous mice exhibit both shortened dendritic arbors and impaired L4>L2/3 synaptic connectivity (Michaelson *et al*., 2018). Dendrites provide a platform for *de novo* synapse formation, suggesting that circuit assembly defects in these mice may arise from impaired neuronal maturation trajectories. It is known that reducing activity in L2/3 sensory cortex neurons during perinatal developmental disrupts their maturation (Cancedda et al., 2007; Gasterstädt et al., 2022), which includes reduced size and complexity of dendrites. Intrinsic membrane excitability (IME) is a cellular feature that gates activity in developing neurons and has been linked to somato-dendritic maturation of developing cortical neurons (Cancedda et al., 2007). Therefore, we hypothesized that *Syngap1* assembles SSC excitatory circuits by regulating IME-dependent maturation of developing glutamatergic neurons. As a first test of this hypothesis, we explored how *Syngap1* expression influences IME in developing L2/3 SSC neurons. Neurons from *Syngap1* heterozygous knockout mice (*Syngap1*^+/-^) were much less intrinsically excitable compared to WT (*Syngap1*^+/+^) controls **(Fig. 1A-C)**. Compared to WT controls, neurons from *Syngap1*^+/-^ mice had dramatically reduced action potential (AP) generation in response to a range of current injections. Consistent with this finding, rheobase was also increased in neurons from *Syngap1*^+/-^ mice, demonstrating that significantly more current is needed to induce a single AP in these neurons compared to WT controls. Importantly, *Syngap1* regulation of IME was cell-autonomous because a similar genotype effect was observed in a sparse population of neurons expressing GFP-Cre in a *Syngap1* conditional knockout (*Syngap1*^+/*fl*^) mouse line **(Fig. 1D-J)**. The effects in *Syngap1*^+/*fl*^ neurons were not due to viral injection or expression of Cre because no phenotypes were observed in *Syngap1*^+l+^ neurons. The effect of *Syngap1* expression on the development of IME was conserved across primary sensory cortex areas in developing mice. Indeed, we observed a similar, albeit less robust *Syngap1* effect in developing visual cortex neurons, which also exhibited reduced membrane excitability (*Fig. S1*). Thus, *Syngap1* expression in developing sensory cortex L2/3 neurons cell autonomously increases IME, which promotes activity in this population during perinatal development.

**Figure 1:**
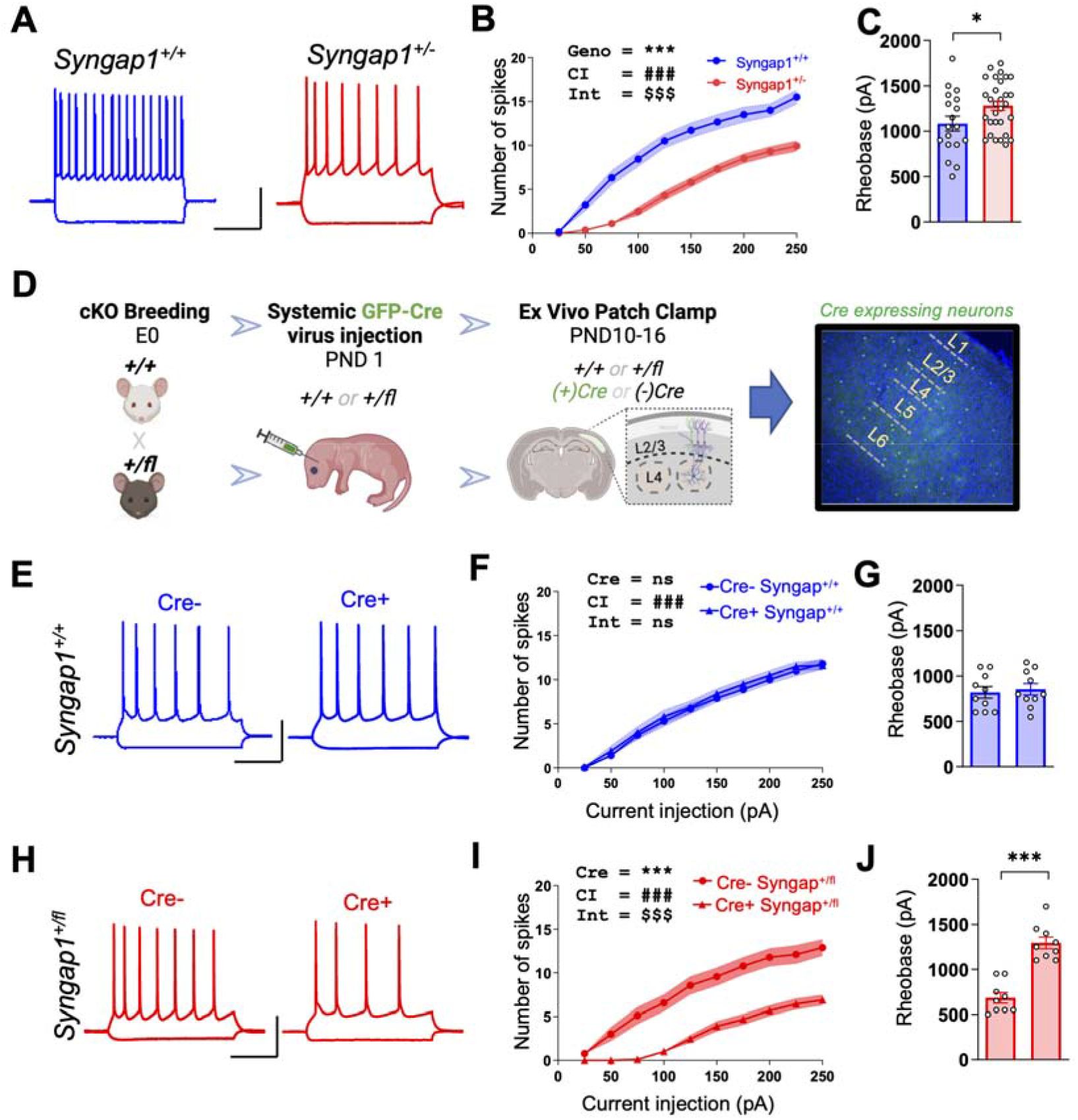
*Syngap1* cell-autonomously regulates regulation of intrinsic membrane excitability in developing SSC L2/3 neurons. (**A**) Representative current clamp traces of AP firing in response to 200 pA current injection from L2/3 SSC excitatory neurons from acute P(10-14) *Syngap1*^+/+^ and *Syngap1*^+/-^ coronal slices (Scale bar 40 mV, 200ms). (**B**) The I/O curve shows reduced AP firing in response to increasing current injections in the Syngap1^+/-^ neurons (Repeated measures ANOVA Genotype F(1.46)=43.71, p<0.0001; Interaction F(9,414)=13.87, p<0.0001, Syngap1^+/+^=18 neurons, 4 mice; Syngap1^+/-^ =30 neurons, 6 mice). (**C**) Bar histogram plots show increased rheobase from Syngap1^+/-^ neurons (Student’s unpaired t test: t (46) =2.171; p=0.0352) in the same set of neurons as in panel (B). (**D**) Schematic showing experimental design of the study involving crossing between Ai9^+/-^ and Syngap1^+/fl^ followed by systemic delivery of r-AAV9-packaged Cre-GFP expressing virus through superior temporal vein to create sparse labelling of neurons with *Syngap1* haploinsufficiency. Acute slices were cut at PND10-14 to measure L2/3 intrinsic excitability. (**E**) Example firing of Cre-GFP negative and Cre-GFP positive L2/3 SSC neurons from *Syngap1*^+/+^ mice. (Scale bar 40 mV, 200 ms). (**F**) Input/Output (IO) curves of SSC L2/3 cells in conditions indicated in panel (E) showing AP firing in response to current injections in L2/3 SSC neurons from *Syngap1*^+/+^ mice with and without Cre expression; RM-Two-way-ANOVA, Cre - F(1,18) = 0.2849, p=0.6000; Injected current x Cre Interaction - F(10, 180) = 0.2713, p=0.9867 from 10 Cre-GFP negative and 10 Cre-GFP positive L2/3 SSC neurons from 3 *Syngap1*^+/+^ mice. (**G**) Bar histogram plots showing rheobase in the same set of neurons as in panel (F); Unpaired t test, t(18)=0.3918, p=0.6998. (**H**) Example firing of Cre-GFP negative and Cre-GFP positive L2/3 SSC neurons from *Syngap1*^+/*fl*^ mice. (Scale bar 40 mV, 200 ms). (**I**) Input/Output (IO) curves of SSC L2/3 cells in conditions indicated in panel (H) showing AP firing in response to current injections in L2/3 SSC neurons from *Syngap1*^+/*fl*^ mice with and without Cre expression; RM-Two-way-ANOVA, Cre F(1, 16) = 28.57, p<0.0001; Interaction - F(10, 16) = 9.740, p<0.0001 from from 9 Cre-GFP negative and 9 Cre-GFP positive L2/3 SSC neurons from 4 Syngap1^+/fl^ mice. (**J**) Bar histogram plots showing rheobase in the same set of neurons as in panel (I); Unpaired t test t(16)=7.042, p=0.0002. Bar graphs represent mean ± SEM, *p < 0.05, **p<0.01, ***p<0.001. For line graphs, symbols equal the population mean, area fill represents SEM. In RM-ANOVA, main effects are denoted by the following symbols: * represents genotype, # represents current injection, and $ represents genotypic and injected current interaction.

What is the cellular mechanism linking cell autonomous *Syngap1* expression to regulation of membrane excitability? Identifying these mechanism(s) is a first step toward exploring potential cause-and-effect relationships between *Syngap1* expression, neuronal IME set points, control of dendritic morphogenesis, and how putative regulation of neuronal maturation directly contributes to building discrete circuit elements during cortical development. IME regulation is heavily linked to function of ion channels that regulate resting membrane potential and membrane resistance (Bando et al., 2022; Smith and Walsh, 2020). We first recorded passive plasma membrane properties from L2/3 SSC neurons within *ex vivo* slices taken from *Syngap1*^+/+^ and *Syngap1*^+/-^ mice shortly after birth (PND1). At this age, action potentials are rudimentary and not present in all neurons (*Fig. S2 A,E*). We observed weak, but significant, effect sizes in RMP (Fig. S2C), rheobase (Fig. S2D), and proportion of neurons that spike (Fig. S2E) in slices taken from newly born *Syngap1* mutants. Non-significant trends were apparent in measures of membrane resistance (Fig. S2B) and potassium conductance (Fig. S2F-G). These results suggested that L2/3 neurons are mildly hypo-excitable in PND~1 *Syngap1* mice. However, by PND~14, weak phenotypes were transformed into large ones, while trends observed at PND~1 emerged as significant. For example, comparing both genotypes revealed that deficient *Syngap1* expression led to clearly hyperpolarized neurons with reduced membrane resistance by the end of the second postnatal week **(Fig. 2A-C)**. This suggested that *Syngap1* deficiency enhances the function of hyperpolarizing ion channel currents. Analysis of action potential (AP) waveforms were consistent with this line of reasoning **(Fig. 2D-I)**. AP depolarization velocity and peak AP amplitude were similar when comparing genotypes **(Fig. 2D,E,F)**. However, the AP repolarization phase was faster **(Fig. 2D,E)**, AP half-width reduced **(Fig. 2D,G)**, while the AP after-hyperpolarization was enhanced **(Fig. 2D,H)**. The AP threshold was also significantly depolarized in mutant neurons compared to controls **(Fig. 2I).** These results suggested that *Syngap1* deficiency leads to enhanced potassium conductance in L2/3 SSC glutamatergic neurons. Direct measurements of potassium currents (*I*_a_) in these neurons supported this hypothesis. Indeed, *Syngap1*^+/-^ neurons exhibited a striking increase in peak *I*_a_ current density **(Fig. 2J,K)**. Plateau density was also changed by *Syngap1* deficiency, but in a different way compared the peak measures, because there was an interaction between genotype and current injection, while there was no main effect on genotype **(Fig. 2J,L)**. Taken together, these data demonstrate that L2/3 neurons in SSC of *Syngap1*^+/-^ mice become progressively less excitable during postnatal development. This timing maps onto the known expression profile of SynGAP proteins in developing mice (Gou *et al*., 2020), which increases sharply in the early postnatal period, and is consistent with the known critical period of this gene (Aceti *et al*., 2015).

**Figure 2:**
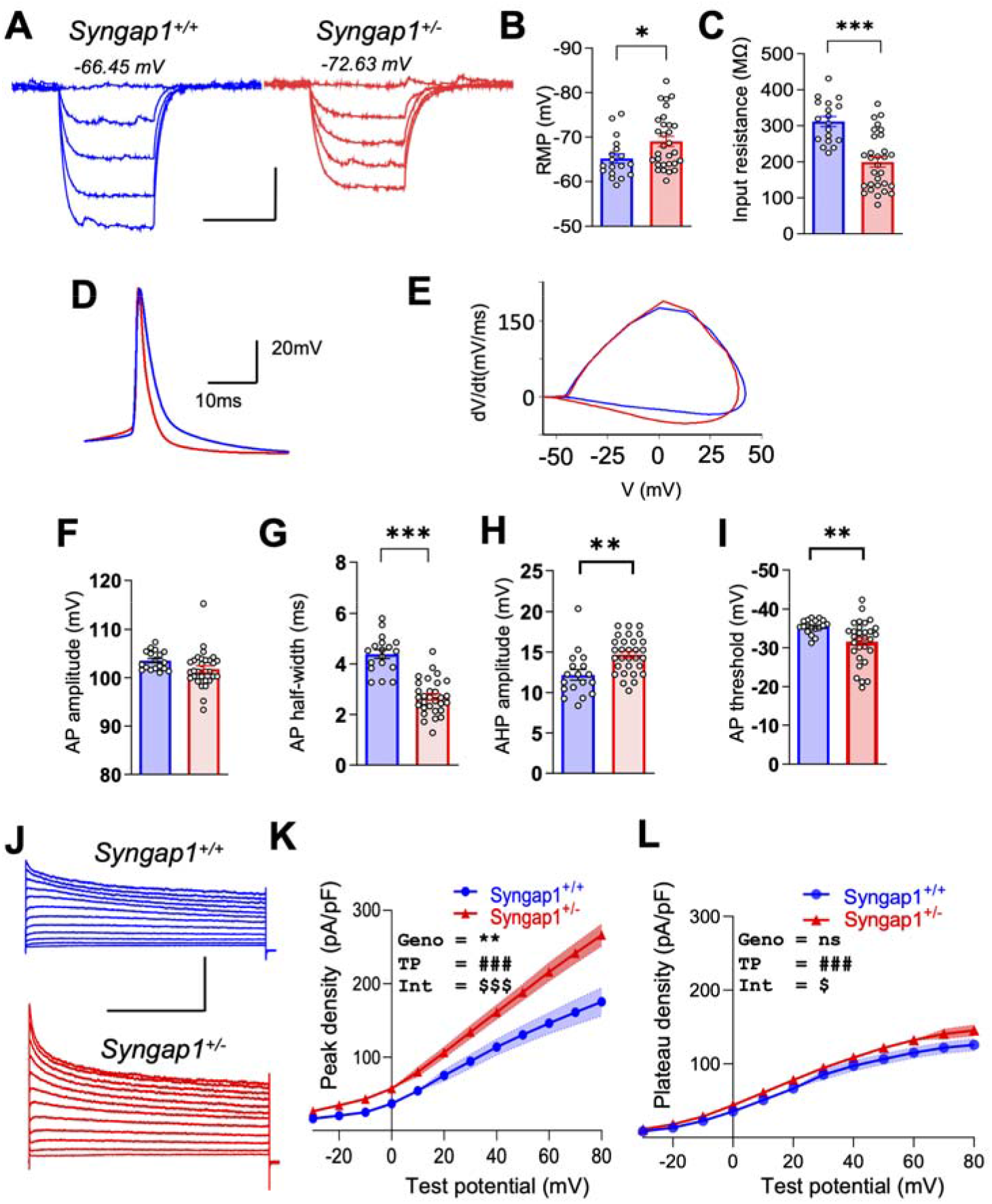
*Syngap1* deficiency alters membrane properties and enhances potassium currents in developing L2/3 SSC pyramidal neurons. (**A**) Representative traces for membrane potential changes in response to hyperpolarizing current steps for neurons from PND10-14 *Syngap1*^+/+^ (blue) or *Syngap1*^+/-^ (red) acute slices (Scale bar 10mV, 100 ms). (**B**) Resting membrane potential was measured for L2/3 SSC neurons (unpaired t-test, t(46)=2.323 p=0.0247; *Syngap1*^+/+^ =18 neurons from 4 mice, *Syngap1*^+/-^ =30 neurons from 6 mice). (**C**) Input resistance from same neuronal populations as in (C); t (46) =5.647, p<0.0001; unpaired t test; *Syngap1*^+/+^ =18 neurons from 4 mice, *Syngap1*^+/-^ =30 neurons from 6 mice. (**D**) Average of AP waveforms generated from 50 pA to 150 pA injections compared across *Syngap1*^+/+^ *and Syngap1*^+/-^ neurons along with representation of (**E**) AP phase plots. (**F**) Plots showing cell-averaged AP amplitudes (unpaired t test, t(46)=1.806, p=0.0774), (**G)** AP halfwidths (unpaired t test, t(46)=8.062, p<0.0001), (**H**) AHP amplitudes (unpaired t test, t(46)=3.434, p=0.0013) and (**I**) AP threshold (unpaired t test, t(46)=2.853, p=0.0065; *Syngap1*^+/+^ =18 neurons from 4 mice, *Syngap1*^+/-^ =30 neurons from 6 mice). (**J**) Representative potassium channel current waveforms in response to 6s long depolarizing pulse of varying test potentials from holding potential of −30mV to +70mV from L2/3 *Syngap1*^+/+^ or *Syngap1*^+/-^ neurons (Scale bar 200 pA, 2s). (**K**) Peak potassium current densities for L2/3 neurons are plotted as a function of test potential; RM 2-way ANOVA, Genotype F (1.29) =8.826, p<0.00133; Test potential x Genotype Interaction F (11,319)=6.664, p<0.0001; Test potential F (11.341)=320.1, p<0.0001; n=15 neurons from 4 *Syngap1*^+/+^ mice and n=18 neurons from 4 *Syngap1*^+/-^ mice. (**L**) Plateau potassium current densities are plotted as a function of test potential; RM 2-way ANOVA, Genotype F (1, 34) = 3.080, p=0.0883; Test potential x Genotype Interaction F (11, 374) = 1.827, p=0.0481; Test potential F (11, 374) = 519.4, p<0.0001; n=15 neurons from 4 *Syngap1*^+/+^ mice and n=18 neurons from 4 *Syngap1*^+/-^ mice. Bar graphs represent mean ± SEM, *p < 0.05, **p<0.01, ***p<0.001. For line graphs, symbols mean neuron means, area fill indicate SEMs, and in RM ANOVAs, main effects are denoted by the following symbols: * represents genotype, # represents test potential and $ represents genotype x test potential interaction.

IME is a highly plastic feature of neurons that is capable of tuning activity so that it remains within a predetermined range, or “set point” (Turrigiano and Nelson, 2004). Neurons self-tune activity by engaging homeostatic plasticity mechanisms that increase or decrease IME, a biophysical process that counteracts sustained decreases or increases in neuronal activity, respectively. *Syngap1* deficiency results in L2/3 neurons with profoundly low IME in development (Fig. 1, 2) that extends into adulthood (Michaelson *et al*., 2018). Given that these neurons are perpetually “stuck” in a low excitability state, we hypothesized that *Syngap1* deficiency impairs homeostatic regulation of IME. To test this idea, cultured slices containing the SSC from P4 *Syngap1*^+/+^ and *Syngap1*^+/-^ mice were prepared and then treated with tetrodotoxin (TTX) or bicuculline (BIC) for at least two days **(Fig. 3A)**. TTX and BIC treatments had the expected impact on neurons from *Syngap1*^+/+^ slices **(Fig. 3B-D, H-J)**, which was to drive an increase and decrease in membrane excitability, respectively. In contrast, in *Syngap1*^+/-^ slices, TTX failed to increase membrane excitability **(Fig. 3E-G)** and BIC failed to reduce it **(Fig. 3K-M),** demonstrating that *Syngap1* deficient L2/3 SSC neurons are unable to regulate membrane excitability in response to persistent changes in activity. Surprisingly, homeostatic regulation of synapse function (i.e., synapse scaling) was present in TTX-treated *Syngap1* deficient neurons, though this process was less robust relative to WT neurons (Fig. S3). Thus, homeostatic plasticity of IME is more sensitive to reduced *Syngap1* expression relative to homeostatic plasticity of synapse function. These studies provide a cellular explanation for perpetually low excitability in *Syngap1* deficient L2/3 neurons – the inability to self-tune IME. They also indicate that *Syngap1* expression maintains neural activity at the “set point” through regulation of IME levels.

**Figure 3:**
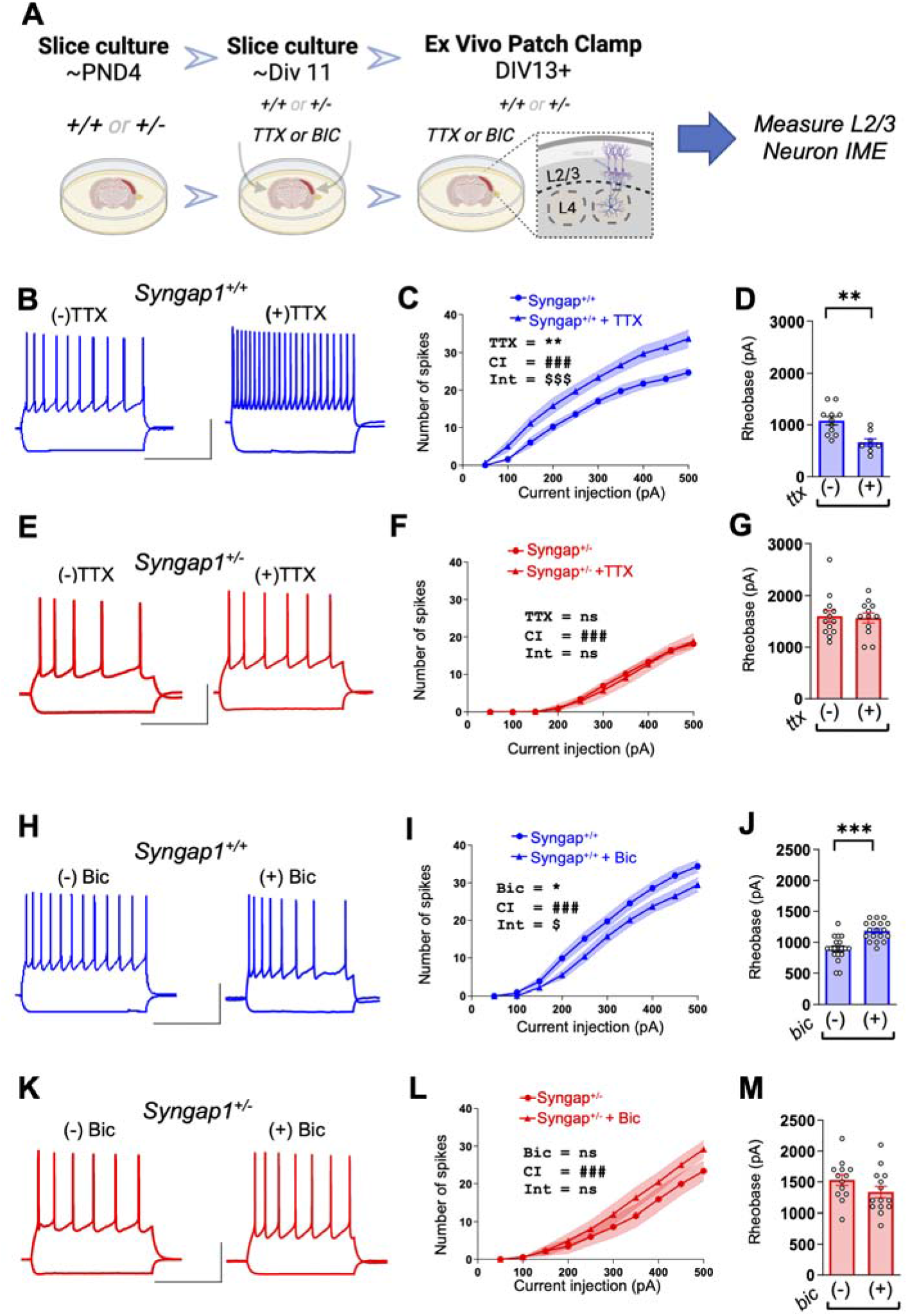
*Syngap1* expression is required for activity dependent tuning of IME in developing SSC L2/3 neurons. **(A)** Schematic timeline of the experimental protocol for organotypic slice culture drug incubations and intrinsic excitability assays for assessing homeostatic plasticity. SSC cortex slices were incubated in a culture medium containing TTX and bicuculline (BIC) for 48 hours to inhibit and excite the neurons chronically. Subsequently, whole cell patch clamp experiments involving intrinsic excitability recordings including AP firing and rheobase were carried out. (**B**) Example spiking of L2/3 SSC pyramidal cells in in response to 200 pA current injection from untreated and TTX treated *Syngap1*^+/+^ (blue) organotypic cultures (Scale bar 40mV, 400ms). (**C**) Input/Output curves of SSC L2/3 cells from untreated and TTX treated *Syngap1*^+/+^ slices (RM 2-way ANOVA; Injected current and TTX Interaction - F (9,189) = 3.494, p=0.0005; Injected current – F(9,189)=229.8, p<0.0001; TTX – F (1,21) = 11.23, p=0.0030; n=12 cells from 4 slices (3 mice) for *Syngap1*^+/+^ vehicle, n=11 cells from 5 slices (3 mice) for *Syngap1*^+/+^ + TTX. (**D**) Rheobase for L2/3 SSC cells from untreated and treated *Syngap1*^+/+^ organotypic cultures (unpaired t test t(17)=3.693, p=0.00188; n=12 cells from 4 slices(3 mice) for *Syngap1*^+/+^ vehicle, n=11 cells from 5 slices (3 mice)for *Syngap1*^+/+^ + TTX). (**E**) Example spiking of L2/3 SSC pyramidal cells in response to 200 pA current injection from untreated and TTX treated *Syngap1*^+/-^ (red) organotypic cultures (Scale bar 40mV, 400ms). (**F**) Input/Output curves of SSC L2/3 cells from untreated and TTX treated *Syngap1*^+/-^ slices (RM 2-way ANOVA: injected current and TTX interaction F (9.207) = 0.2929, p=0.9761; TTX - F (1,23) = 0.03811, p=0.8469; Injected current – F (9,207) = 150.5, p<0.0001; n=13 cells from 5 slices (3 mice) for *Syngap1*^+/-^ + vehicle and n=12 cells from 4 slices (2 mice) for *Syngap1*^+/-^ + TTX). (**G**) Rheobase for vehicle and TTX for *Syngap1*^+/-^ mice (unpaired t test, t(23)=0.2183, n=13 cells from 5 slices (3 mice) for *Syngap1*^+/-^ + vehicle and n=12 cells from 4 slices (2 mice) for *Syngap1*^+/-^ + TTX). **(H)** Example firing of neurons in response to 200 pA current injections from untreated and Bicuculline treated *Syngap1*^+/+^ organotypic cultures of mouse SSC (Scale bar 40mV, 400ms). **(I)** Input/Output curves of SSC L2/3 cells from untreated and bicuculline treated *Syngap1*^+/+^ organotypic cultures [RM-ANOVA, Injected current x Bicuculline Interaction – F (9,315) = 2.420, p<0.0114; Injected current - F(9,315) = 362.6, p<0.0001 Bicuculline – F (1,35) = 4.419, p=0.0428; n=19 cells from 6 slices (3 mice) for *Syngap1*^+/+^ + vehicle, n=18 cells from 5 slices (3 mice) for *Syngap1*^+/+^ + Bicuculline. (**J**) Rheobase measures in L2/3 SSC neurons in untreated and bicuculline treated organotypic slices from *Syngap1*^+/+^ mice [unpaired t test, t(35)=4.962, p<0.0001; n=19 cells from 6 slices (3 mice) for *Syngap1*^+/+^ + vehicle, n=18 cells from 5 slices (3 mice) for *Syngap1*^+/+^ + Bicuculline. (**K**) Example firing of neurons in response to 200 pA current injections from untreated and Bicuculline treated *Syngap1*^+/-^ organotypic cultures of mouse SSC (Scale bar 40mV, 400ms). **(L)** Input/Output curves of SSC L2/3 cells from untreated and bicuculline treated *Syngap1*^+/-^ organotypic cultures [RM 2 way-ANOVA, Injected current x Bicuculline Interaction - F(9,225) = 1.703, p<0.0894; Injected current - F(9,225) = 114, p<0.0001 Bicuculline - F(1,25) = 1.057, p=0.3138; n=13 cells from 5 slices (2 mice) for *Syngap1*^+/-^ + vehicle, n=14 cells from 4 slices (2 mice) for *Syngap1*^+/-^ + Bicuculline. (**M**) Rheobase measures in L2/3 SSC neurons in untreated and bicuculline treated organotypic slices from *Syngap1*^+/-^ mice [unpaired t test, t(25)=1.503, p<0.1455; n=13 cells from 5 slices (2 mice) for *Syngap1*^+/-^ + vehicle, n=14 cells from 4 slices (2 mice) for *Syngap1*^+/-^ + Bicuculline. Bar graphs represent mean ± SEM, *p < 0.05, **p<0.01, ***p<0.001. For line graphs, symbols mean neuron means, area fill indicate SEMs, and in RM ANOVAs, main effects are denoted by the following symbols: * represents TTX/Bicuculline treatment, # represents current injection and $ represents TTX/Bicuculline treatment x injected current interaction.

*Syngap1*, as the name implies, is a well-established genetic regulator of dendritic spine signaling that directly controls excitatory synapse structure, function, and plasticity (Gamache et al., 2020; Kilinc et al., 2018). Thus, it is possible that changes in developmental IME in *Syngap1* deficient neurons is a consequence of its known functions within excitatory synapses. To test this idea, we measured homeostatic plasticity mechanisms in a mouse model expressing mutations in an alternatively spliced *Syngap1* isoform known to regulate synapse-specific mechanisms. Indeed, we and others have shown that the α1 c-terminal protein variant of *Syngap1* is highly enriched in dendritic spine synapses where it functions to regulate biology within this compartment (Araki et al., 2020; Gou et al., 2020; Kilinc et al., 2022). Synaptic compartmentalization and biological functions of α1 is imparted by expression of a PDZ binding motif (PBM) present within an alternatively spliced terminal exon. We recently reported the construction and characterization of a mouse line with an α1 C-terminal isoform-specific PDZ domain binding motif (PBM) mutation (Kilinc et al., 2022). This mutation disrupts SynGAP-α1 proteins from binding to PDZ proteins at the synapse, but importantly does not alter total SynGAP protein expression. The consequence of altered PBM binding of this isoform is impaired dendritic spine Ras/Erk signaling, enhanced L2/3 SSC neuron baseline excitatory synapse function, and weakened Hebbian-like plasticity of excitatory synapses (Araki et al., 2020; Kilinc et al., 2022). If *Syngap1* regulates IME through biological functions originating within excitatory synapses, then IME plasticity may be impaired in L2/3 neurons from PBM mice. To test this idea, we treated cultured slices from WT or *Syngap1*-PBM mice with TTX and then measured how these treatments changed IME **(Fig. 4A)**. TTX treatment enhanced membrane excitability in WT neurons as expected **(Fig. 4A-C)**. Surprisingly, membrane excitability was also enhanced in response to TTX treatment in *Syngap1-PBM* neurons **(Fig. 4D-F),** demonstrating that homeostatic regulation of IME is intact in neurons lacking fully functional α1 isoforms. To assess homeostatic plasticity of excitatory synapses in this line, we next performed the TTX treatment in cultured *Syngap1-PBM* slices and measured mEPSC properties. As expected, TTX treatment enhanced mEPSC amplitude in WT neurons, reflecting engagement of homeostatic plasticity at glutamatergic synapses **(Fig. 4G-H)**. In contrast, TTX treatment failed to alter mEPSC properties in *Syngap1-PBM* neurons **(Fig. 4J-K)**, indicating that functionally impaired α1 isoforms disrupt homeostatic synapse plasticity. Intact IME homeostatic plasticity combined with deficient homeostatic synapse plasticity in PBM mice demonstrates that *Syngap1*-mediated IME phenotypes are dissociated from synapse phenotypes. This marks the discovery of a novel *Syngap1* gene function – direct control of IME - that may be related to protein functions outside dendritic spine synapses.

**Figure 4:**
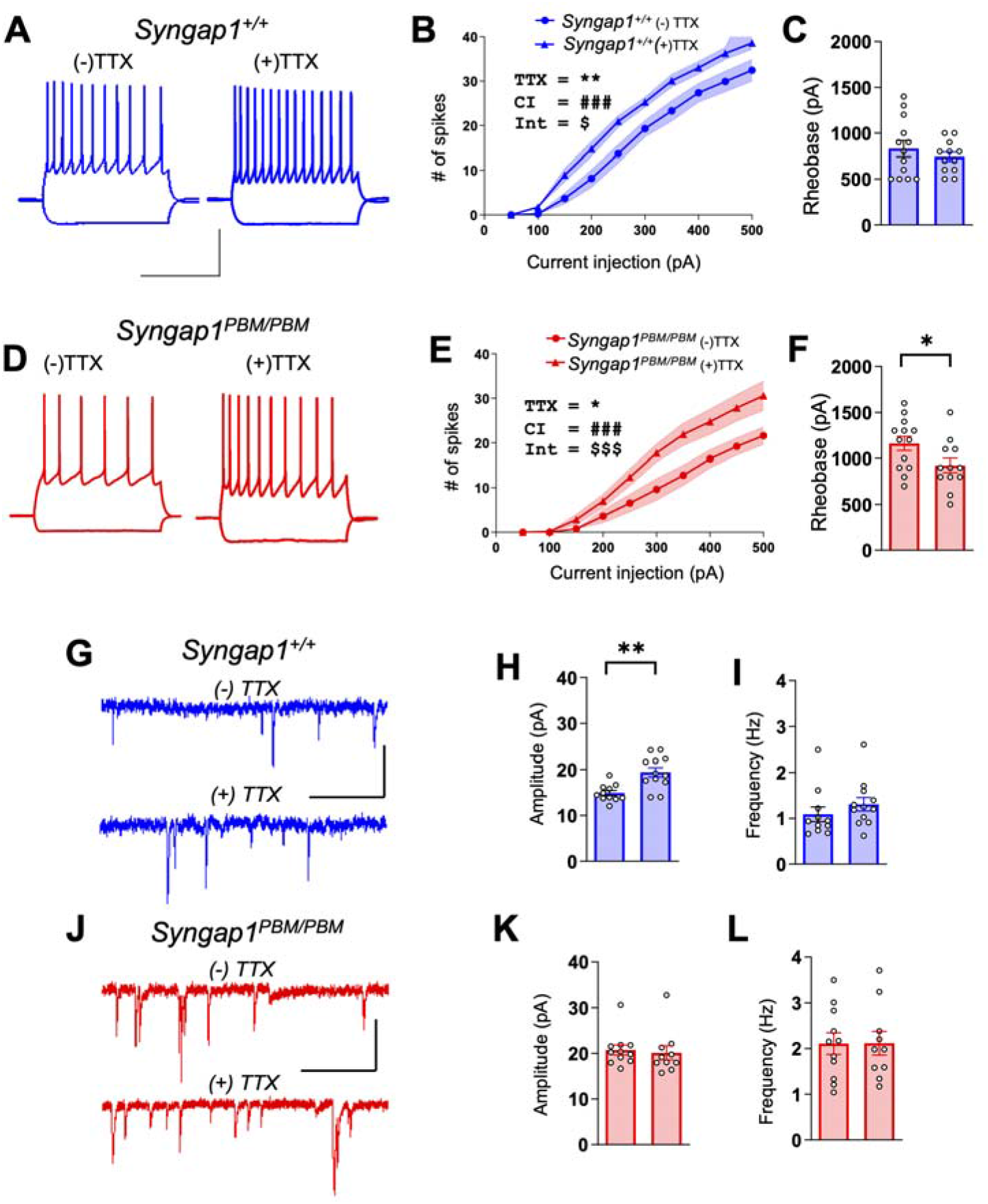
The canonical dendritic spine isoform, SynGAP-a1, regulates homeostatic synapse plasticity but not membrane excitability. (**A**) Example spiking of L2/3 pyramidal cells in SSC in response to 200 pA current injection from untreated and TTX treated *Syngap1*^+/+^ organotypic cultures (Scale bar 40mV, 400ms). (**B**) Input/ Output curves of SSC L2/3 cells from untreated and TTX treated *Syngap1*^+/+^ organotypic cultures (Repeated measures 2-way ANOVA Injected current x TTX Interaction - F (9,207) = 2.133, p=0.0282; Injected current - F (9, 207) = 257.9, p<0.0001 TTX - F (1,23) = 9.521, p=0.0052; n=13 cells from 6 slices (4 mice) for *Syngap1*^+/+^ *vehicle*, n=12 cells from 5 slices (3 mice) for *Syngap1*^+/+^ + TTX. **(C)** Rheobase for vehicle and TTX for *Syngap1*^+/+^ mice (unpaired t test t(23)=0.8555, p=0.4011; n=13 cells from 6 slices (4 mice) for *Syngap1*^+/+^ vehicle, n=12 cells from 5 slices (3 mice) for *Syngap1*^+/+^ + TTX). (**D**) Example spiking of L2/3 SSC pyramidal cells in response to 200 pA current injection from untreated and TTX treated *Syngap1^PBM/PBM^* organotypic cultures (Scale bar 40mV, 400ms). (**E**) Input/ Output curves of SSC L2/3 cells from untreated and TTX treated *Syngap1^PBM/PBM^* organotypic cultures (RM 2-way ANOVA Injected current x TTX Interaction - F (9,207) = 4.247, p<0.0001; Injected current - F (9, 207) = 119.6, p<0.0001 TTX - F (1,23) = 6.730, p=0.0162; n=13 cells from 7 slices (5 mice) for *Syngap1^PBM/PBM^*, n=12 cells from 6 slices (4 mice) for *Syngap1^PBM/PBM^* + TTX. (**F**) Rheobase for L2/3 SSC cells from untreated and treated *Syngap1^PBM/PBM^* organotypic cultures (unpaired t test t(23)=2.008, p=0.0421; n=13 cells from 7 slices(5 mice) for *Syngap1^PBM/PBM^* vehicle, n=12 cells from 6 slices (4 mice) for *Syngap1^PBM/PBM^* + TTX). (**G**) Representative mEPSC responses from L2/3 SSC cells in organotypic cultures from untreated and TTX (1 μM) treated *Syngap1*^+/+^ organotypic cultures (Scale bar 40 pA, 1s). (**H**) Quantification of mEPSC amplitudes for L2/3 SSC cells in *Syngap1*^+/+^ organotypic cultures subjected to each condition in (G) (unpaired t test, t (21) = 3.808, p=0.0010; n=11 cells from 4 slice (2 mice) for *Syngap1*^+/+^, n=12 cells from 5 slices (3 mice) for *Syngap1*^+/+^ + TTX). (**I**) Quantification of frequency of mEPSCs for L2/3 SSC cells in organotypic cultures subjected to each condition in (G) (Mann Whitney’s exact test, U=41, p=0.1335; n=11 cells from 4 slice (2 mice) for *Syngap1*^+/+^, n=12 cells from 5 slices (3 mice) for *Syngap1*^+/+^ + TTX). (**J**) Representative mEPSC responses from L2/3 SSC cells from untreated and TTX (1 μM) treated *Syngap1^PBM/PBM^* organotypic cultures (Scale bar 40 pA, 1s). (**K**) Quantification of mEPSC amplitudes for L2/3 SSC cells in *Syngap1^PBM/PBM^* organotypic cultures subjected to each condition in (J) (Mann Whitney exact test, U = 41.5, p=0.3589; n=11 cells from 4 slices (3 mice) for *Syngap1^PBM/PBM^*, n=10 cells from 5 slices (3 mice) *Syngap1^PBM/PBM^* + TTX) (**L**) Quantification of frequency of mEPSCs for L2/3 SSC cells in *Syngap1^PBM/PBM^* organotypic cultures subjected to each condition in (J) (unpaired t test, t(19)=0.0305, p=0.9760; n=11 cells from 4 slices (3 mice) for *Syngap1^PBM/PBM^* n=10 cells from 5 slices (3 mice) for *Syngap^PBM/PBM^* + TTX). Bar graphs represent mean ± SEM, *p < 0.05, **p<0.01, ***p<0.001. For line graphs, symbols mean neuron means, area fill indicate SEMs, and in RM ANOVAs, main effects are denoted by the following symbols: * represents TTX treatment, # represents current injection and $ represents TTX treatment x injected current interaction.

We hypothesized that *Syngap1* regulation of IME may contribute to activity-dependent maturation of neurons that instructs assembly of cortical feed-forward circuits. This hypothesis emerged from the observation that adult *Syngap1* mice have small L2/3 neurons with shortened dendritic arbors and impaired excitatory connectivity in SSC (Michaelson et al., 2018). However, these animals express spontaneous absence-like seizures that worsen throughout adulthood (Ozkan et al., 2014; Sullivan et al., 2020). Thus, it is unknown if these circuit phenotypes emerge from pathology related to growing excitability imbalances during adulthood or from abnormally developing L2/3 SSC neurons. Therefore, we traced dendritic arbors from developing SSC L2/3 neurons in *Syngap1*^+/+^ and *Syngap1*^+/-^ mice (Fig. S4A). Dendritic arbors from developing neurons in *Syngap1*^+/-^ mice were less extensive and less complex compared to neurons from *Syngap1*^+/+^ animals **(Fig. 5A-C;** Fig. S4B-C). Shortened dendrites observed in mutant animals also had lower spine density (Fig. S4B). Combined, these dendrite and spine phenotypes suggested that developing neurons with *Syngap1* deficiency express fewer anatomical excitatory connections. We next determined if these anatomical features translated into functional synaptic phenotypes. *m*EPSC analysis was next performed **(Fig. 5D)**, which revealed a significant reduction in frequency **(Fig. 5E)**, but not amplitude **(Fig. 5F)**, of L2/3 SSC neurons. This finding was consistent with reduced excitatory connectivity. To directly measure functional excitatory connectivity within a defined circuit motif, we stimulated L4 inputs onto SSC L2/3 neurons **(Fig. 5G)**. This experiment revealed an impairment in feed-forward excitatory connectivity **(Fig. 5H-I)** within a circuit required for cortical processing of touch (Sachidhanandam et al., 2013). Altered dendritic morphogenesis in mutant animals was not due to compensation of an abnormally functioning cortex. Sparse Cre expression within a conditional *Syngap1* mouse line (*Syngap1*^+/*fl*^ mice) revealed a nearly identical dendritic phenotype to that of *Syngap1*^+/-^ mice **(Fig. 5J-K)**. Moreover, we also observed a reduction in dendritic arborization in L2/3 primary visual cortex (V1) neurons from *Syngap1*^+/-^ mice (Fig. S4C-D). Impaired IME is correlated with altered dendritic maturation in *Syngap1*^+/-^ mice (Figs. 1 and 3). Given that our hypothesis states that *Syngap1* control of IME regulates dendritic maturation, and that IME plasticity was intact in *Syngap1* PBM mice (Fig. 4), we suspected that neurons from these mice would fail to express the arrested maturation phenotype. Indeed, neurons from *Syngap1-PBM* mice had dendritic trees that were similar to WT controls **(Fig. 5L-N)**. These results underscore a strong correlation between how *Syngap1* regulates IME and how L2/3 SSC neurons mature during perinatal development.

**Figure 5:**
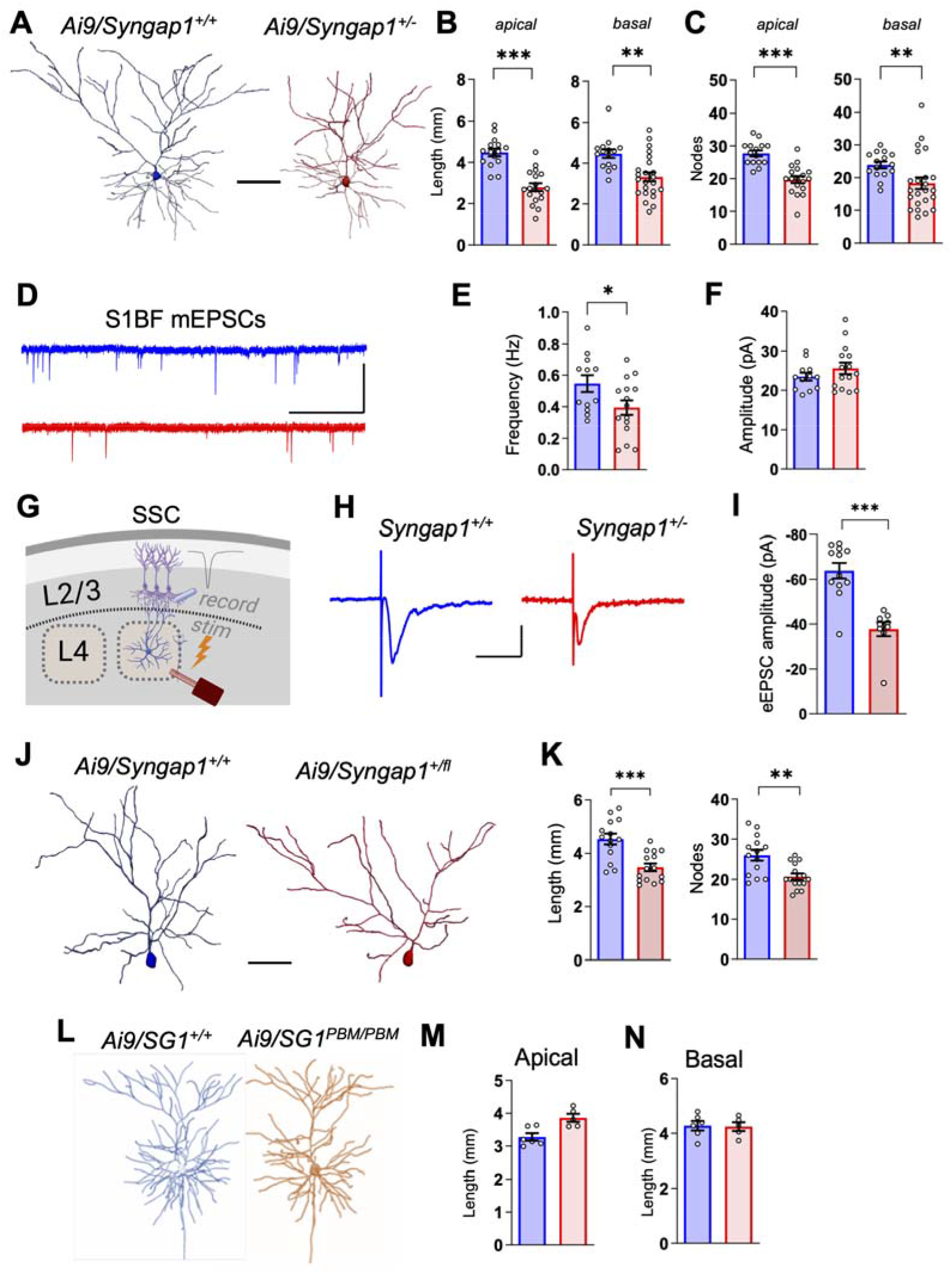
*Syngap1* cell-autonomously regulates developmental dendritic morphogenesis in *L2/3* SSC glutamatergic neurons. (**A**) SSC L2/3 dendrite reconstructions from developing *Syngap1*^+/+^ (Blue) and *Syngap1*^+/-^ (Red) mice expressing the Ai9 allele (Scale bar, 100um). (**B**) Bar histogram graphs depicting the total length of both apical (total length - unpaired t-Test t(31)=6.288, p<0.0001; *Syngap1*^+/+^ = 3 mice, 15 neurons; *Syngap1*^+/-^ = 3 mice, 18 neurons) and basal (total length - unpaired t-Test; t(36)=3.568, p = 0.0010; *Syngap1*^+/+^ = 4 mice, 15 neurons; *Syngap1*^+/-^ = 5 mice, 23 neurons) dendritic arbors. (**C**) Bar histograms depicting the # of nodes of apical (# of nodes - unpaired t-Test; t(31)=5.625, p<0.0001; *Syngap1*^+/+^ = 3 mice, 15 neurons; *Syngap1*^+/-^ = 3 mice, 18 neurons) and basal (# of nodes – Mann-Whitney’s exact test: U=82.50, p = 0.0062; *Syngap1*^+/+^ = 4 mice, 15 neurons; *Syngap1*^+/-^ = 5 mice, 23 neurons) dendritic arbors. (**D**) Representative traces for mEPSCs from PND10-14 *Syngap1*^+/+^ and *Syngap1*^+/-^ L2/3 SSC neurons (Scale bar = 4s, 40pA). (**E-F**) Bar histogram plots for frequency (E; Unpaired t test: t (25)=2.160, p=0.0406, *Syngap1*^+/+^ = 3 mice, 12 neurons; *Syngap1*^+/-^ =4 mice, 15 neurons) and amplitude (F; unpaired t test, t(25)=1.124, p=0.2715, *Syngap1*^+/+^ = 3 mice, 12 neurons; *Syngap1*^+/-^ =4 mice, 15 cells) of mEPSCs in L2/3 SSC neurons. (**G**) Cartoon depicting experimental setup for measuring L4>L2/3 feed-forward excitation (FFE) in barrel region of SSC. (**H**) Representative traces depicting L2/3 excitatory neuron, L4-evoked eEPSCs from acute *Syngap1*^+/+^ and *Syngap1*^+/-^ PND10-14 thalamocortical slices (Scale bar 20 pA, 50 ms). **(I)** Bar histogram plot of eEPSC amplitude in L2/3 neurons (Mann-Whitney’s exact test: U= 8, p=0.0005); *Syngap1*^+/+^ = 4 mice, 12 neurons; *Syngap1*^+/-^ = 3 mice, 9 neurons. (**J**) Examples of reconstructed L2/3 neurons from SSC of *Syngap1*^+/+^ and *Syngap1*^+/*fl*^ mice (Scale bar, 100μm). **(K**) Bar histogram plots depicting apical length and # of apical nodes (total length - unpaired t-Test; t(27)=4.269, p = 0.0002; # of nodes-unpaired t test; t(27)=3.518, p = 0.0016; *Syngap1*^+/+^ = 4 mice, 14 neurons; *Syngap1*^+/fl^ = 4 mice, 15 neurons). (**L**) Examples of reconstructed L2/3 neurons from SSC of *Syngap1*^+/+^ and *Syngap1^PBM/PBM^* mice (Scale bar, 100μm). (**M**-**N**) Bar histogram graphs depicting the total length of both apical (total length – Mann Whitney’s exact test U=4, p=0.0519; *Syngap1*^+/+^ = 2 mice, 6 neurons; *Syngap1^PBM/PBM^* = 2 mice, 5 neurons) and basal (total length – Mann Whitney’s exact test; U=19, p = 0.9999; *Syngap1*^+/+^ = 2 mice, 6 neurons; *Syngap1^PBM/PBM^* = 2 mice, 5 neurons) dendritic arbors. Values represent neuron means; error bars indicate SEMs. *p < 0.05, **p<0.01, ***p<0.001.

Identification of the cellular mechanism(s) that connect *Syngap1* expression to IME regulation would enable testing of a potential cause-and-effect relationship between IME, L2/3 maturation, and associated circuit assembly during cortical development. Membrane properties of *Syngap1*^+/-^ mice indicated that this gene constrains potassium conductance in developing L2/3 SSC glutamatergic neurons and that this process contributes to aspects of membrane excitability (Fig. 2). Therefore, we hypothesized that lowering elevated potassium channel function in neurons from *Syngap1*+/- mice would normalize IME to WT neuron levels. To do this, we sought to express a dominant negative form of the Kv4.2 potassium channel subunit, *dn*Kv4.2 (Yuan et al., 2005), in developing *Syngap1* mice. We selected this molecular tool for two reasons. First, Kv4 potassium channels substantially contribute to *I*_a_ currents in developing sensory cortex neurons and are dominant in repolarizing the membrane during AP firing (Carrasquillo et al., 2012; Norris et al., 2010; Pathak et al., 2016). Indeed, *I*_a_ currents were elevated and AP repolarization was accelerated in *Syngap1* deficient neurons (Fig. 2). Second, expression of *dn*Kv4.2 has been shown to increase IME by suppressing peak *I_a_* currents in sensory cortex glutamatergic neurons (Yuan et al., 2005). *Syngap1*^+/+^ and *Syngap1*^+/-^ littermates were transfected with mRuby and *dn*Kv4.2 at E15.5, preparation of acute slices containing the SSC occurred at PND12-16, and then mRuby (+) or (-) neurons were patch clamped in slices from each genotype **(Fig. 6A-B).** In these studies, we obtained four electrophysiological readouts that reflect changes in IME. Application of two-way ANOVAs to each data set enabled comparisons across transfection state and genotype. Comparing genotypes in untransfected neurons revealed a clear decrease in membrane excitability in developing *Syngap1*^+/-^ neurons relative to *Syngap1*^+/+^ controls, as there was an effect of genotype in all four measures **(Fig. 6D-G).** This represents a further replication of the principle finding in this study, which is that *Syngap1* expression regulates IME in developing cortical neurons. Importantly, *dn*Kv4.2 expression changed each measure from neurons from both genotypes in a manner consistent with increased IME, an expected result because it is known to reduce potassium channel function in pyramidal neurons. However, the effects of *dn*KV4.2 were more pronounced in *Syngap1*^+/-^ neurons compared to *Syngap1*^+/+^ controls. For example, an interaction was detected between genotype and transfection in the spiking I/O measures, and the effect size was clearly larger in *Syngap1*^+/-^ neurons compared to *Syngap1*^+/+^ neurons **(Fig. 6D)**. In two of the other three measures, including rheobase and input resistance, there was significance noted in post-hoc comparisons between (+) and (-) transfection conditions in *Syngap1*^+/-^ neurons, but not in the *Syngap1*^+/+^ populations **(Fig. 6E,F,G)**. Given that there were main effects of genotype and transfection condition in these two measures, but no detected interactions, these outcomes support an interpretation that the effect of transfection state on *Syngap1*^+/+^ neurons was less robust compared to *Syngap1*^+/-^. Finally, post-hoc comparisons in all four measures revealed that there was no difference between untransfected *Syngap1*^+/+^ neurons and transfected *Syngap1*^+/-^ neurons. In each case, these populations were essentially overlapping, and the means were nearly identical (grey arrows; **Fig. 6D-G)**. Thus, *dn*Kv4.2 expression in developing *Syngap1*^+/-^ neurons returns basic measures of IME back to WT levels.

**Figure 6:**
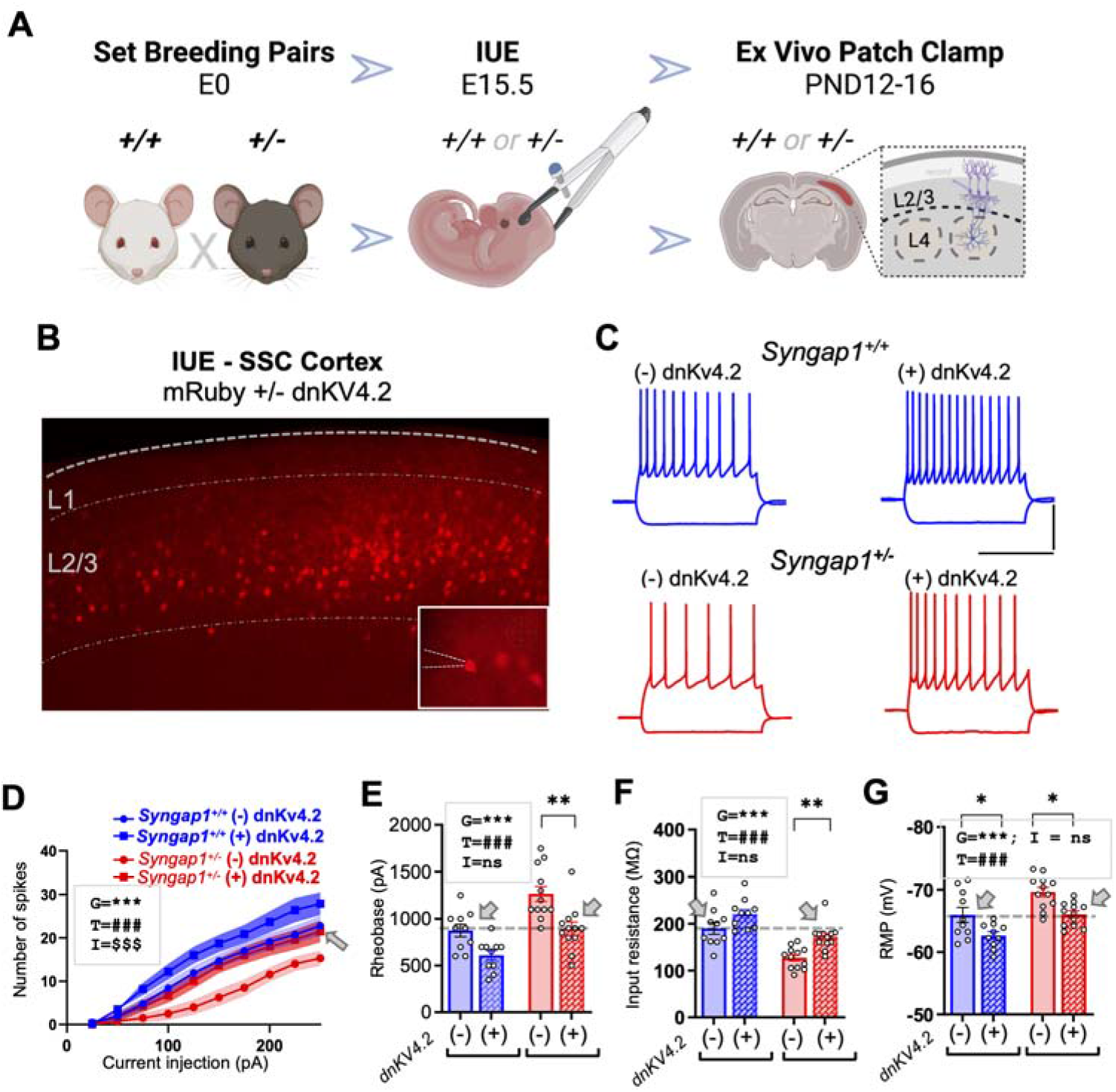
Disrupting potassium channel function in *Syngap1*^+/-^ L2/3 SSC neurons normalizes membrane excitability. (**A**) Schematic showing experimental timeline for in utero electroporation targeting of mRuby and dnKv4.2 plasmid in SSC L2/3 neurons at E15.5 followed up by whole cell patch clamp experiments performed at P12-16. (**B**) Confocal image an acute P14 brain slice showing dnKv4.2 and mRuby plasmids expressed in L2/3 SSC neurons following E15.5 IUE. Inset shows whole cell patch of a transfected neuron (**C**) Representative current clamp traces of AP firing in response to 100 pA current injection from untransfected dnKv4.2(-) and transfected dnKv4.2(+) L2/3 excitatory neurons from PND12-14 *Syngap1*^+/+^ and *Syngap1*^+/-^ mice (Scale bar 40mV, 400ms). (**D**) I/O curve for AP firing in response to increasing current injections in dnKv4.2(-) or dnKv4.2(+) neurons from acute slices prepared from *Syngap1*^+/+^ or *Syngap1*^+/-^ mice (RM 2-way ANOVA, Interaction - F(27, 360) = 4.667, p<0.0001; Genotype - F(3, 40) = 7.096, p=0.0006; Injected current - F(40, 360) = 25.74, p<0.0001; *Syngap1*^+/+^ = 10 pairs of (-)dnKv4.2 and (+)dnKv4.2 neurons, Syngap1+/- = 12 pairs of (-)dnKv4.2 and (+)dnKv4.2 neurons). (**E**) Rheobase measurements in these populations; Two-way ANOVA, Interaction - F(1, 40) = 0.5672, p=0.4558; Genotype - F(1, 40) = 23.24, p<0.0001; dnKv4.2 treatment - F(1, 40) = 20.47, p<0.0001; *Syngap1*^+/+^ = 10 pairs of (-)dnKv4.2 and (+)dnKv4.2 neurons, *Syngap1*^+/-^ = 12 pairs of (-)dnKv4.2 and (+)dnKv4.2 neurons. (**F**) Input resistance in these populations; Two-way ANOVA, Interaction - F(1, 40) = 0.5603, p=0.4585; Genotype - F(1, 40) = 36.36, p<0.0001; dnKv4.2 treatment - F(1, 40) = 16.68, p=0.0002; *Syngap1*^+/+^ = 10 pairs of (-)dnKv4.2and (+)dnKv4.2neurons, *Syngap1*+/- = 12 pairs of (-)dnKv4.2 and (+)dnKv4.2 neurons. (**G**) Resting membrane potential in these neurons; Two-way ANOVA, Interaction - F(1, 40) = 0.03109, p=0.8609; Genotype - F(1, 40) = 18.40, p=0.0001; dnKv4.2 treatment - F(1, 40) = 19.60, p<0.0001; *Syngap1*+/+ = 10 pairs of (-)dnKv4.2 and (+)dnKv4.2 neurons, *Syngap1*^+/-^ = 12 pairs of (-)dnKv4.2 and (+)dnKv4.2 neurons. Bar graphs represent mean ± SEM, *p < 0.05, **p<0.01, ***p<0.001. For line graphs, symbols mean neuron means, area fill indicate SEMs, and in two-way ANOVAs, main effects are denoted by the following symbols: * represents genotype, # represents dnKv4.2 treatment and $ represents genotype x dnKv4.2 treatment interaction. Arrows represent baseline measurements of (-)dnKv4.2 in *Syngap1*^+/+^ mice.

If *Syngap1* regulation of neural activity through control of IME is a mechanism that drives somatodendritic maturation during the perinatal period, then readjusting low IME in developing *Syngap1*^+/-^ L2/3 neurons to WT-like levels through *dn*Kv4.2 expression should enhance this measure of maturation. To test this idea, we measured how *dn*Kv4.2 expression in L2/3 neurons influenced dendritic morphogenesis during early postnatal development. This was performed by infusing a cell tracer in either transfected or untransfected neurons from acute slices prepared from *Syngap1*^+/+^ or *Syngap1*^+/-^ mice (Fig. 7A-C; Fig. S5). Expression of *dn*Kv4.2 in *Syngap1*^+/-^ neurons significantly enhanced dendritic arborization compared to untransfected controls of the same genotype (Fig. 7C-D). This finding revealed a mechanistic link between *Syngap1*-mediated control of IME and dendritic morphogenesis occurring during the E16 > PND10+ window of development. However, it remained unknown to what extent alterations in dendritic morphogenesis from *Syngap1* deficient neurons directly contribute to excitatory feed-forward circuit formation in developing sensory cortex. To resolve this issue, we patch clamped neurons in L2/3 from *Syngap1*^+/-^ mice that were either positive or negative for *dn*Kv4.2 and then performed assays of functional connectivity in SSC. Indeed, expression of *dn*Kv4.2 significantly increased *m*EPSC frequency in transfected neurons compared to untransfected controls (Fig. 7E). There were no observed effects of *dn*Kv4.2 on *m*EPSC amplitude. To determine how targeting potassium channel hyperfunction in *Syngap1*^+/-^ neurons directly influenced translaminar feed-forward connectivity in SSC, we measured L4-evoked excitatory synaptic currents from L2/3 neurons (Fig. 7G). *dn*KV4.2 expression in neurons from mutant mice had significantly enhanced evoked excitatory synaptic function in this circuit (Fig. 7H), a finding consistent with elevated mEPSC frequency. In marked contrast, *dn*Kv4.2 expression in neurons from *Syngap1*^+/+^ mice had no effect on L2/3 pyramidal cell somato-dendritic maturation or excitatory synapse connectivity (Fig. S5A-G). Thus, L2/3 dendritic maturation and subsequent integration into feed-forward excitatory circuits requires *Syngap1* control of IME during the perinatal period of development.

**Figure 7:**
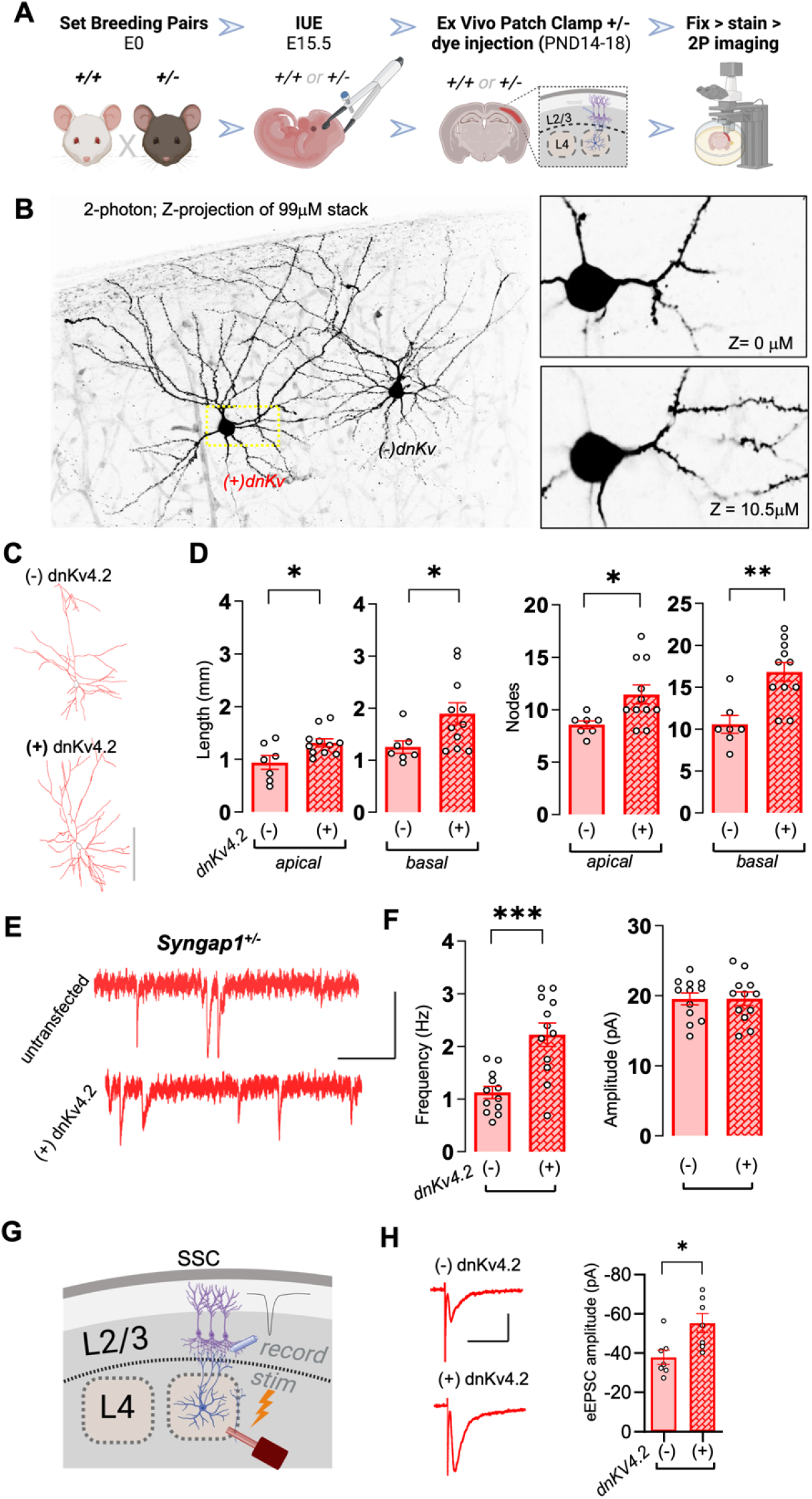
Enhancing IME in *Syngap1*^+/-^ SSC neurons accelerate dendritic morphogenesis and promotes excitatory circuit connectivity. (**A**) Schematic showing experimental strategy and timeline for crossing of *Syngap1*^+/-^ males with Syngap1^+/+^ females followed by in utero electroporation of embryos at E15.5 to target mRuby w/ dnKv4.2 in L2/3 SSC neurons for electrophysiological and morphological measurements. (**B**) Two photon maximum intensity projection image of Neurobiotin-filled streptavidin-Alexa 488 labelled untransfected (-) dnKv4.2 and transfected (+) dnKv4.2 neurons from acute slices prepared from PND14 *Syngap1*^+/-^ mice. **(C)** Representative reconstructions of (-) dnKv4.2 and (+) dnKv4.2 L2/3 SSC neurons from acute coronal slices prepared from *Syngap1*^+/-^ mice; scale bar 100 μm. (**D**) Quantification of apical and basal dendritic length and nodes of traced (-) dnKv4.2 and (+) dnKv4.2 neurons from *Syngap1*^+/-^ mice (Apical dendrites, total length – Mann Whitney’s exact test U=16, p=0.0441; # of nodes-Mann Whitney’s exact test U=12, p=0.0108; Basal dendrites, total length – Mann Whitney’s exact test U=12, p=0.0154; # of nodes Mann Whitney’s exact test t test, U=7, p=0.0024; *Syngap1*^+/-^ = 2 mice, (-) dnKv4.2 = 7 neurons, (+) dnKv4.2 = 11 neurons). (**E**) Representative mEPSCs traces from (-) dnKv4.2 untransfected and (+) dnKv4.2 transfected L2/3 SSC neurons from PND12-14 *Syngap1*^+/-^ acute slices; Scale bar 1s, 40pA. (**F**) *Left* - Bar histograms showing quantification of mEPSCs frequencies from (-) dnKv4.2 untransfected and (+) dnKv4.2 transfected L2/3 SSC neurons from PND12-14 *Syngap1*^+/-^ mice (unpaired t test t (22) =4.394; p=0.0002). **(F)** *Right* - mEPSC amplitudes from same neurons (unpaired t test t (22) =0.01888, p=0.9851). **(G)** Cartoon depicting experimental setup for measuring feed-forward excitation from L4>L2/3 excitatory neurons. **(H)** *Left* - representative L4-evoked L2/3 EPSC traces from untransfected or transfected L2/3 excitatory neurons from acute *Syngap1*^+/-^ slices, Scale bar 20 pA, 50 ms. **(H)** *Right* - Bar histogram plot of evoked EPSC amplitudes in these populations (Mann Whitney’s exact test, U= 6, p=0.0175). Bar graphs represent mean ± SEM, *p < 0.05, **p<0.01, ***p<0.001.

## Discussion

Our results provide mechanistic insight into how gene expression during neuronal development promotes the assembly of *in vivo* excitatory cortical circuit motifs. We elucidated a mechanism for how an individual gene shapes activity-dependent assembly of a translaminar feed-forward circuit through IME-dependent regulation of neuronal maturation in the post-migration phase of cortical development **(Fig. S6)**. In its most simplistic form, IME can be viewed as a collection of voltage-gated sodium, potassium, calcium, and voltage-independent leak channels on the plasma membrane **(Fig. S6A)**, which act in concert to regulate how much synaptic current is required to depolarize neurons and induce AP firing (Moody and Bosma, 2005; Vacher et al., 2008). Therefore, IME acts as a biophysical bridge that connects thousands of synaptic inputs to somatic AP firing. This is a remarkably important neuronal feature because pathologically low IME could effectively negate the action of all synaptic inputs, whereas high IME could result in AP firing from the spurious activity of just one or two synaptic inputs. We find that a principal function of *Syngap1* expression within L2/3 sensory cortex neurons is to tune IME to levels that ensure neuronal activity reaches a threshold that induces dendritic maturation during this critical period of perinatal cortical development **(Fig. S6A)**. In this model, IME dynamically regulates neuronal activity over a protracted temporal window, from days to weeks around the perinatal period, which corresponds to the period of extensive cortical axodendritic differentiation, synaptogenesis, and subsequent circuit assembly. Neuronal maturation itself can influence IME levels (van der Velden et al., 2012; van Elburg and van Ooyen, 2010), suggesting that IME changes in *Syngap1* deficiency could be a consequence of altered pace of neuronal development. However, we demonstrate that re-establishing WT-like levels of IME in *Syngap1* deficient L2/3 neurons unleashed neuronal maturation (Figs. 6 and 7), including enhanced dendritic morphogenesis, which in-turn promoted translaminar feed-forward excitatory circuit formation **(Fig. S6B;** Fig. 7). This supports the idea that dendritic maturation defects in *Syngap1* deficient neurons is caused by reduced IME, rather than vice-versa. Moreover, prior studies have shown that activity within sensory cortex L2/3 glutamatergic neurons during this developmental period is a signal that promotes somato-dendritic maturation (Cancedda et al., 2007; Gasterstädt et al., 2022). Genetic lowering of IME in WT neurons through exogenous expression of a potassium channel subunit, or direct disruptions to neuronal spiking in developing sensory cortex neurons using a chemogenetic approach, both resulted in an arrested dendritic maturation phenotype that resembled observations in *Syngap1* deficient neurons.

It has been shown that regulating IME in cortical progenitors can alter migration dynamics (Bando et al., 2014; Vitali et al., 2018), which can control their ultimate final position within the cortex. This finding raises the possibility that *Syngap1* regulation of neural progenitors may explain our maturation phenotypes. However, sparse induction of *Syngap1* deficiency after birth, a period in development when cortical neurons have finished migrating and all fates have been specified, was sufficient to induce both impaired IME (Fig. 1) and somato-dendritic maturation (Fig. 5). Importantly, these results do not preclude a role for *Syngap1* in neurogenesis or neuronal migration. Rather, they demonstrate that regulating *Syngap1* expression after this phase of cortical development is sufficient to impair a form of neuronal maturation that is defined by post-migration somato-dendritic differentiation, which provides a platform to assemble upper lamina sensory cortex circuits. Any effects of *Syngap1* expression on neurogenesis or patterning would in principle interact with the post-migratory maturation mechanisms described in this study, leading to a complex and multidimensional impact on circuit structure/function.

The finding that *Syngap1* regulates IME-mediated somato-dendritic development in cortical neurons was unexpected because this gene is a well-known and potent regulator of excitatory synapse maturation and plasticity (Gamache et al., 2020; Kilinc et al., 2018). Indeed, *Syngap1* has been shown to regulate the maturation of excitatory synapses in developing sensory cortex excitatory neurons (Clement et al., 2013). Moreover, it was recently shown to regulate experience-dependent plasticity of glutamatergic neuron ensembles in L2/3 SSC, as well as activity-dependent plasticity of excitatory synaptic inputs on these neurons (Llamosas et al., 2021). Because studies of *Syngap1* have largely focused on deciphering its unique functions at the synapse, theories that seek to explain how it regulates neural functions are synapse-centric (Gamache et al., 2020; Walkup et al., 2016). While *Syngap1* is indeed a potent regulator of excitatory synapse biology, and that these functions contribute to the development of neural functions (Kilinc et al., 2022), our current findings extend the known function of the *Syngap1* gene beyond the synapse and into the somato-dendritic compartment. Our current findings demonstrate that this gene functions to control IME through regulation, at least in part, of ion channel currents on the somato-dendritic membrane. This view is supported by the effect of *Syngap1* deficiency on potassium currents in L2/3 neurons (Fig. 2). As a result, these new findings demonstrate that this gene is crucial for assembling nascent circuits by controlling the rate of activity-dependent dendritic maturation through setting appropriate levels of membrane excitability. Thus, *Syngap1* expression during the mouse perinatal critical period of cortical development regulates the two major neuronal processes for building mature circuits in sensory cortex: IME regulation of somatodendritic differentiation that aids assembly (this study), and regulation of excitatory synapse plasticity for experience-dependent refinement of circuits (Llamosas et al., 2021). Experiments using the PBM mouse line suggest that these two distinct cellular functions of the *Syngap1* are dissociable. The PBM mouse line harbors SNVs in the *Syngap1* gene that selectivity disrupts the function of the α1 isoform by interfering with binding to PDZ proteins (Kilinc et al., 2022). These SNVs were shown to have no impact on the other SynGAP C-terminal variants. The α1 isoform has specialized functions compared to the other C-terminal SynGAP protein variants, β, α2, and γ, and that these specialized functions are restricted to excitatory synapses. It has been established by multiple groups that α1 is more concentrated in dendritic spines compared to the other isoforms (Araki et al., 2020; Gou et al., 2020). There, it controls GTPase signaling at excitatory synapses that regulates dynamic changes in synaptic strength. Consistent with this, homozygous PBM mice, which express α1 protein completely devoid of PDZ binding, exhibit altered maturation of excitatory synapses on L2/3 SSC pyramidal neurons, impaired LTP, and express behaviors that reflect disrupted cognitive processing (Kilinc et al., 2022). However, in this current study, altered α1 isoforms had no effect on homeostatic plasticity of IME (but did regulate homeostatic plasticity – Fig. 4), which was in stark contrast to homeostatic plasticity regulation in *Syngap1* Heterozygous KO mice, which express a ~50% reduction in all protein isoforms (Kilinc et al., 2022). These findings indicate that one or more of the other C-terminal variants contribute to IME regulation. This is consistent with what is known about their spatial-temporal expression profiles, where they are less enriched in spines and more enriched in the somato-dendritic compartment (Araki et al., 2020; Gou et al., 2020), the locus of ion channels that most directly regulate IME. Furthermore, dendritic morphogenesis was not arrested in neurons from the PBM line, which is consistent with the finding of normal IME plasticity in response to chronic changes in activity. The combined effect of *Syngap1* gene function on synaptic and intrinsic components of L2/3 sensory cortex neuron excitability provide a new framework for understanding the emergence of multidimensional pathological phenotypes in the human population.

It was somewhat surprising to find that *dn*KV4.2 expression and subsequent elevation of IME in WT neurons had no effect on dendritic morphology or translaminar synaptic connectivity (Fig. S5). Because reduced activity in the perinatal period leads to impaired neuronal maturation and disrupted circuit assembly in L2/3 neurons, one might predict that enhancing activity by raising IME would do the opposite. The most likely explanation for these results is that maturation mechanisms induced by activity reach saturation once a critical threshold of activity is achieved, with additional activity failing to drive further differentiation. In *Syngap1* mutants, reduced IME results in low activity levels. Thus, these neurons likely do not receive sufficient activity during the perinatal period to saturate molecular signaling pathways that drive morphogenesis. Based on this rationale, our data strongly implicate *Syngap1* as a regulator of activity “set-points” within individual L2/3 neurons. This theory posits that a persistent change to a neuron’s activity level engages homeostatic cellular mechanisms to drive neuronal activity in the opposing direction to push levels back toward the intrinsic “set point” (Turrigiano and Nelson, 2004). Indeed, *Syngap1* heterozygosity prevented plasticity of IME induced by either tetrodotoxin or bicuculline (Fig. 3). Thus, these neurons lost a powerful mechanism for driving activity back to the intrinsic “set point”, leaving them “stuck” in a state of low excitability. This is supported by the prior finding of reduced IME in these same neurons in adult *Syngap1* mice (Michaelson et al., 2018).

Regulation of IME by major NDD genes may be a generalizable mechanism that instructs activity-dependent assembly of nascent cortical circuits. Indeed, IME is regulated by other highly penetrant NDD genes. *FMR1/Fmr1*, which is linked to Fragile X syndrome (FXS), and *SHANK3/Shank3*, which is linked to Phelan-McDermid syndrome, have also been shown to regulate membrane excitability. *Shank3* expresses proteins that are present at the PSD, where they link NMDAR function to calcium influx related to AMAPR trafficking required for Hebbian forms of synapse plasticity (Delling and Boeckers, 2021). However, *Shank3* also regulates the function of voltage-gated ion channels, such as HCN1 channels, which form I_*h*_ current in neurons. I_*h*_ currents are important for regulating membrane excitability and dysregulation of I_*h*_ currents in reduced culture preparations were linked to altered dendritic morphogenesis (Yi et al., 2016). Shank3 is also important for regulation of homeostatic plasticity of membrane excitability (Tatavarty et al., 2020). However, it remains unresolved how Shank3 expression regulates assembly of defined circuits *in vivo. Fmr1* regulates synaptic protein synthesis/trafficking, protein/protein interactions, and controls neuronal excitability (Contractor et al., 2015). *Fmr1* KO mice recapitulate many symptoms of the clinical population including heightened neuronal excitability. The contributing mechanisms are continually being resolved, however, altered ion channel regulation/function are thought to participate (Deng et al., 2021). Interestingly, there are conflicting reports as to the effect that loss of *Fmr1* has on some ion channels. For example, some studies show a reduction for A-type K^+^ channels (*I_KA_*) (Gross et al., 2011), while others show an increase (Lee et al., 2011) in expression/function. Furthermore, I_h_ has been observed to be elevated in the hippocampus (Brager et al., 2012), while down-regulated in L5B of SSC (Zhang et al., 2014). Clearly, *Fmr1* has divergent effects on various ion channels, not only between but within brain regions depending on cell-type. This highlights the importance of studying the relationships between ion channels, neuronal excitability, and neuronal maturation on a circuit-by-circuit and region-by-region manner. This was the motivation behind this current study, which was to study these relationships in a specific neuronal subtype (L2/3 pyramidal neurons) that is a component of a defined circuit motif (translaminar feed-forward excitatory connectivity), which is known to regulate a neural process (sensory processing) with implications for behavioral control. Therapeutic interventions aimed at selective targeting of neurons with developmentally impaired IME may serve to prevent various forms of circuit connectivity alterations by correcting dendritic maturation.

## Acknowledgments

We thank Jeanne Nerbonne for sharing the dnKV4.2 plasmid. This work was supported in part by NIH grants from the National Institute of Mental Health (MH096847 and MH108408 to G.R. and MH105400 to C.A.M.), the National Institute for Neurological Disorders and Stroke (NS064079 to G.R.) and the National Institute for Drug Abuse (DA034116 and DA036376 to C.A.M.). VA and SDM were supported by training fellowships from Leon and Friends and SynGAP Research Fund, respectively.

## Author Contributions

V.A. and S.D.M. performed experiments, designed experiments, analyzed data, and co-wrote the manuscript. M.A. and M.K. performed experiments, designed experiments, and analyzed data. C.A.M. designed experiments, interpreted data, and edited the manuscript. G.R. conceived the study, designed experiments, interpreted data, co-wrote the manuscript, and edited the manuscript.

## Declaration of Interests

The authors declare no competing financial interests.

## Materials and Methods

### Mouse strains and breeding strategies

All animal procedures were conducted in accordance with the National Institutes of Health Guide for the Care and Use of Laboratory Animals and protocols were approved by the Scripps Research Institute and UF-Scripps Biomedical Research Animal Care and Use Committees. The design and validation of constitutive *Syngap1* KO (*Syngap1*^+/-^) (Kim et al., 2003), conditional *Syngap1* KO (*Syngap1*^+/*fl*^) (*Clement et al., 2012*), and *Syngap1*^PBM/PBM^ (Kilinc et al., 2022) lines have been described previously. The Ai9 Cre reporter strain (#007905) was purchased from Jackson Laboratories. The Ai9 allele was heterozygous in resulting offspring from crosses with the various *Syngap1* lines. All experimental mice were used before weaning, where they were housed with the mother, on a 12-hour normal light-dark cycle.

#### Cell autonomous studies

For determining cell autonomous effects of *Syngap1* on various electrophysiological phenotypes, P1 *Syngap1*^+/*fl*^ mice were injected with an rAAV9-packaged Cre-GFP expressing virus (pENN.AAV.CamKII.HI.GFP-Cre.WPRE.SV40: Addgene #105551-AAV9: Titer = 5.66×10^13^ GC/ml) via the superficial temporal vein (STV) of PND1 mouse pups as described previously (Ozkan et al., 2015). Sparsely labelled neurons (~1-5% of total) expressed GFP when Cre was present which led to *Syngap1* haploinsufficiency in <u>*Syngap1*^+/*fl*^</u> mice. Furthermore, for cell autonomous effects of *Syngap1* on dendritic morphology, P1 *Syngap1*^+/*fl*^ crossed to *Ai9* mice were injected with an rAAV9-packaged Cre-expressing virus (pENN.AAV.hSyn.Cre.WPRE.hGH; Addgene # 105553-AAV9; Titer 2.3 x 10^13^). Briefly, pups were sedated by covering them with ice for 3 min. STV was visualized using a handheld transilluminator (WeeSight; Respironics), and a pair of standard reading glasses. Virus solution was prepared by 1:50 dilution of stock solution (final ~10^12 GC/mL) in Dulbecco’s phosphate-buffered saline (PBS), also supplemented with 0.001% pluronic-F68. Virus solution (50 nL) was injected using a 100 nL Nanofill syringe attached with a 34-gauge Nanofill beveled needle (World Precision Instruments). A correct injection was verified by noting blanching of the vein. After the injection, pups were returned to the incubator until active and then returned to their dam until future studies.

### Acute ex vivo brain slice preparation

Acute coronal (IME and mEPSC assays) or thalamocortical slices (eEPSC assays), from PND10-14 mice were cut using standard methods as previously described (Clement et al., 2013; Llamosas et al., 2020; Michaelson *et al*., 2018). Briefly, mice were decapitated, and brains rapidly removed in ice-cold cutting solution containing (in mM): 119 NaCl, 2.5 KCl, 26.3 NaHCO3, 2.5 CaCl2, 1 NaH2PO4, 1.3 MgSO4, 10 Ascorbic acid and 11 D-Glucose, and equilibrated with 95% O2 and 5% CO2 (pH 7.4, ~300 mOsm). Brain slices containing the somatosensory cortex (SSC) or visual cortex (V1) were cut (~350 μm) using a vibrating microtome (Candem Instruments) before placing into a submersion chamber containing warmed (32-34°C) artificial cerebrospinal fluid (aCSF), composed of (mM): 125 NaCl, 2.5 KCl, 24 NaHCO3, 2 CaCl2, 1.25 NaH2PO4, 2 MgSO4, and 10 D-Glucose, and equilibrated with 95 % O2 and 5 % CO2 (pH 7.4, ~300 mOsm) for 30-40 minutes. Following this, slices were maintained in bubbled aCSF at room temperature until used for recordings (1-6 h).

### Whole cell patch clamp electrophysiology of acute ex vivo brain slices

Whole-cell patch clamp experiments were conducted at room temperature from visually identified L2/3 neurons in SSC or V1 using infrared DIC optics and regular spiking was confirmed in current clamp. Recordings were made using borosilicate glass pipettes (3-6 MΩ; 0.6 mm inner diameter; 1.2 mm outer diameter; Harvard Apparatus). All signals were amplified using Multiclamp 700B (Molecular Devices), filtered at 2 KHz, digitized at 10 KHz, and stored on a personal computer for off-line analysis. Analog to digital conversion was performed using the Digidata 1440A system (Molecular Devices). Data acquisitions and analyses were performed using pClamp software packages (Clampex (10.7) and Clampfit (11.6) respectively; Molecular Devices).

For current clamp and evoked excitatory postsynaptic current (eEPSC) recordings, the following internal solution was used (in mM): 130 potassium gluconate, 5 KCl, 10 HEPES, 0.25 EGTA, 10 phosphocreatine disodium, 0.5 Na-GTP and 4 Mg-ATP (pH 7.3, 285-290 mOsm). For miniature excitatory postsynaptic current (mEPSC) recordings, the following internal solution was used (in mM): 120 CsCl, 5 NaCl, 1 MgCl2, 10 HEPES, 10 EGTA, 3 Mg-ATP and 0.3 Na-GTP (pH 7.3, 285-290 mOsm). Cells with access resistance >30 MΩ or were unstable (>20 % change in this measure) were discarded from further analysis. Cells included in the study had resting membrane potentials corrected post hoc for liquid junction potential (measured at 10mV).

Evoked EPSC (eEPSC) experiments were performed as previously described (Michaelson *et al*., 2018). Briefly, slices were incubated with 100 μM picrotoxin. L4 was stimulated by placing a concentric bipolar stimulating electrode (FHC, 25 μm inner diameter; 125 μm outer diameter) in the center of a visually defined barrel by transillumination at 4x. Neurons were recorded directly above (vertical to) the stimulus site in L2/3 ~50-150uM from L1. The stimulation intensity (0.2 ms, constant-current pulses) was regulated by a stimulus isolation unit (ISO-Flex, A.M.P.I). A minimum stimulus intensity protocol was employed (6 sweeps with 15 s ISI) in current clamp, as eEPSPs appeared to be more stable than eEPSCs recorded in voltage clamp. The stimulus intensity was gradually increased until the emergence of an eEPSP with a 30-50 % failure rate. This stimulus intensity was used in voltage-clamp mode to elicit eEPSCs from ~15 sweeps (15 s ISI). eEPSCs were quantified by averaging the peak amplitudes, within a 30 ms post-stimulus window, excluding failures. Only one neuron was recorded from a single barrel column and a maximum of 2 neurons per slice. mEPSCs were recorded in TTX (1 μM), picrotoxin (100 μM) and APV (100 μM) at a holding potential of −75 mV. For each recording, ~5 minutes of data were acquired and subsequently analyzed using Mini Analysis software (Synaptosoft Inc).

Examining IME properties was conducted in current clamp mode using a number of established protocols. To determine spike numbers, a family of depolarizing current injections (500 ms, 25-50 pA increments) from SSC and V1 L2/3 neurons was employed. Rheobase was measured by depolarizing steps of 50 pA for 300 msec and then computing the current injection which elicit the first action potential. Resting membrane potential (Vm) was measured at I=0. Membrane voltage changes were measured in response to hyperpolarizing current steps whose slope of *V/I* curve was calculated to determine the input resistance (R_input_).

For AP waveform analysis, action potential analysis function in Clampfit 11.6 was used. AP waveforms generated from 50 pA to 150 pA injections were selected for evaluation. Measurements of AP threshold, AP halfwidth, AHP amplitude and AP amplitude were chosen for analysis. Individual action potential measurements were averaged for a given neuron. AP phase plots were generated in Clampfit 11.6 by plotting the first derivative of the AP voltage against AP voltage.

For potassium current recordings, voltage-gated outward K^+^ currents were recorded from L2/3 SSC neurons using 1μM TTX and 100 μM CdCl_2_ in the bath solution to block Na^+^ and Ca^2+^ currents, respectively. K channel current waveforms were measured in response to 6 s long depolarizing pulses of varying test potentials in voltage clamp mode from holding potentials of −30mV to +70mV from L2/3 SSC neurons. Peak and plateau K^+^ current densities for L2/3 neurons were plotted as a function of test potential.

For PND 0-1 cortical plate neuron recordings, the brain was dissected to obtain cerebral cortices in icecold postnatal ACSF (Bahrey and Moody 2003, Beier and Barish 2000) bubbled with carbogen. 200 μm coronal slices were cut, followed by incubation for an hour in postACSF (which contained (in mM) 115 NaCl, 4.3 KCl, 2CaCl2, 2 MgCl2, 0.28 MgSO4, 0.22 KH2PO4, 0.85 Na2HPO4, 27 NaHCO3, and 25 glucose for 60–90 min before recording). Higher resistances of 8-10 MΩ were pulled and filled with potassium internal solution, which contained (in mM) 110 K Gluconate, 5 glucose, 1.1 CaCl2, 6 KCl, 5 MgCl2, 40 HEPES, 3 Mg-ATP, and 0.5 Na-GTP, pH to 7.25, osmolarity 290 mosm to obtain whole cell patch clamp recordings in both voltage and current clamp from pyramidal cells.

### Organotypic culture preparation

Organotypic cultures were prepared from PND4-6 *Syngap1*^+/-^ and *Syngap1*-PBM mice using standard protocols described elsewhere (Michaelson et al., 2021). In brief, mice were decapitated, and brains quickly removed in a laminar fume hood under sterile conditions. Brains were immersed in ice cold slicing solution [consisting of Hank’s balanced salt solution (HBSS) +0.6% D-glucose and 30 μM Kynurenic acid]. Approximately four ~350 μM coronal slices (per hemisphere) containing SSC were cut using a vibratting microtome (Candem Instruments). Slices were allowed to rest in fresh slicing solution at 4 °C for 30-45 minutes before being mounted on individual semiporous membrane inserts (2 slices per insert) placed in 6 well plates with 1.1mL of culture media [50% Minimal Essential Medium (MEM; Gibco), 25% heat-inactivated horse serum (Gibco), 25% HBSS(Thermofisher Scientific), supplemented with: 1 mM Glutamax, 1% D-glucose (Sigma-Aldrich), 0.5 mM L-ascorbic acid (Sigma Aldrich), and 25 U/mL penicillin/streptomycin (Sigma-Aldrich)]. Slices were maintained at 37°C in a 5%/95% (CO_2_/air) humidified incubator. Media was changed every other day for the duration of the experiment. For homeostatic plasticity experiments, slices were treated with either 1 μM TTX or 25 μM Bicuculline at DIV11 for 48 hours followed by whole cell patch clamp electrophysiology.

### Morphological analysis of in vivo labelled L2/3 neurons

For dendritic tracing, PND1 *Syngap1*^+/-^, *Syngap1^PBM/PBM^*, or *Syngap1*^+/*fl*^ expressing the *Ai9* Cre reporter allele were injected with a rAAV9-packaged Cre-expressing virus (<u>pENN.AAV.hSyn.Cre.WPRE.hGH; Addgene # 105553-AAV9; Titer 2.3 x 10^13^)</u>, via STV injection described previously. At PND14, animals were deeply anesthetized with pentobarbital (Nembutal) and transcardially perfused with 4% PFA/PBS (wt/vol). Extracted brains were postfixed in 4% PFA/PBS at 4°C for 10 h and cryoprotected in 20% sucrose/PBS (wt/vol) at 4°C for 24 h. Brains were cut on a vibratome (500 μm thickness) collecting SSC. Slices were placed in 6-well plates and submerged in ScaleA2 solution, at 4°C, and gentle shaking to create transparent tissue. Slices were mounted for confocal imaging. Brain weight was measured at different time points: pre-ScaleA2 (T0) and post-ScaleA2 (T24h-T5weeks), and the linear expansion (E) was calculated by comparing each post-ScaleA2 measures with the pre-ScaleA2 baseline. The cube root of pre/post ScaleA2 comparison results in estimating the time-fold linear expansion. Consequently, for comparing morphological parameters of both genotypes, the estimation of the extent of linear expansion promoted by ScaleA2 solution was necessary. Appling the coefficient of expansion (CoE = 1/E), the measures addressing the total length of dendritic branches were rescaled accordingly. Due to all confocal sessions being conducted within the third and fifth weeks post/ScaleA2, re-scaling CoE applied was calculated by averaging each CoE from the third to the fifth week during ScaleA2 treatment. The formula [1 – (1/E)] was used to determine the percentage of linear expansion in all groups.

### Confocal Imaging for genetic labeling/morphology experiments

Tissue was considered for imaging after 2 weeks in the ScaleA2 solution. Brain slices were mounted in Petri dishes, covered in agarose, as described (Hama et al., 2011). Three-dimensional image stacks were collected (x: 2048, y: 2048 pxl; step size 1.00 μm) using an Olympus FV1000 laser-scanning confocal microscope equipped with a water immersion objective lens (ULTRA 25x, NA 1.05, Olympus). A computer-based tracing system (Neurolucida360; MicroBrightField) was used to generate threedimensional neuron tracings that were subsequently visualized and analyzed with NeuroExplorer (MicroBrightField) to quantify the total dendritic length and number of branch points. Neurons were selected based on the following criteria: 1) The soma was present in the middle of the stack (~150 μm ± 30 μm) to ensure the accurate reconstruction of an entire dendritic arbor; 2) the neuron was distinct from other neurons to allow for identification of branches; 3) the neuron was not truncated in any obvious way. As the clearing method enhanced the quality of our images, spine density was determined using the same set of images previously acquired for the tracing experiment. As previously described (Aceti et al., 2015), ten to fifteen dendritic segments of SSC L2/3 neurons (20–90 μm in length) at P14 mice were collected and considered for analysis. All measurements were performed by an experimenter blind to the experimental conditions. Pictures were visualized and elaborated with Neurolucida 360 software (MicroBrightField).

### In utero electroporation

We followed standard protocols for this procedure with minor modifications (Au - Matsui et al., 2011). 7-week-old or greater female C57/BL6j strain (Jackson Labs #000664) were mated with *Syngap1*^+/-^ male mice and plugs were tracked to create timed pregnancies. E15.5 embryos within timed pregnant dams were electroporated with electrode placement meant to target the SSC. Briefly, dams were first anesthetized with isoflurane (5% induction, 2.5 % maintenance). After fixation on a prewarmed surgery platform and sterilization of the skin with Betadine, the abdominal cavity was opened through 2 incisions along the midline to expose the uterine horns. Embryos were removed and injected one at a time with 2 μL (2μg/μl) DNA containing pAAV-CAG-mRuby2 and CMV-dnKv4.2 using pulled glass micropipettes (WPI) into the lateral ventricle. For better visibility and location of the injection site, 0.1% Fast Green (Sigma) was added to the DNA solution. Targeted labeling of neuronal progenitors residing in the SSC, destined to be L2/3 neurons, was performed by applying electric pulses (45 V; 50 ms on; 950 ms off, 5 pulses) with an electric stimulator (Cuy21EDIT, Bex C. Ltd.) using round electrodes (CUY650P5, CUY650P7, Nepa Gene) encapsulating the embryonic head. Uterine horns with embryos were nudged gently back into the abdominal cavity, which was then filled with prewarmed PBS containing carprofen diluted at 0.5 mg/ml (final dose 5mg/kg) and antibiotics (Penicillin-Streptomycin 1:100). The abdominal muscle was sutured using absorbable sutures and the skin was closed using non-absorbable Ethilon nylon sutures (Ethicon). Animals were allowed to recover on a heating pad before replacement to their home cages. mRuby (Addgene #99123; pAAV-CAG-mRuby2) and CMV-dnKv4.2 plasmids (Barry et al., 1998) were prepared using a standard endotoxin-free maxiprep and additionally purified by ethanol precipitation. cDNA preps were resuspended to ~2 mg/ml in dH20.

### Labeling and imaging of neurons from acute *ex vivo* slices

SSC neurons from PND14 *Syngap1*^+/+^ and *Syngap1*^+/-^ mice (untransfected/transfected with dnKv4.2 via in utero electroporation) were filled during whole cell patch clamp recordings of IME with the following internal solution: 126 K-gluconate, 4 KCl, 10 HEPES, 5 MgATP, 0.3 NaGTP, 1 EGTA, 0.3 CaCl2 and 0.2% neurobiotin (pH 7.27, 290 mOsm/L). Neurons from *ex vivo* slices were then stained, imaged, reconstructed, and subjected to the same morphologic analyses as previously described (Michaelson et al., 2020). Briefly, acute slices were fixed with 4% PFA (Thermo Fisher Scientific) for 24 h at 4°C, then transferred to 0.2% sodium azide in PBS for subsequent storage overnight at 4°C. Slices were washed for 3 × 10 min periods in PBS. For blocking and permeabilization, slices were kept at room temperature in wells containing 0.3% Triton X-100 in PBS and 4% normal goat serum (Sigma-Aldrich) for 3 hours. Slices were then incubated overnight with streptavidin conjugated AlexaFluor-488 (1:1000; Invitrogen) in 0.3% Triton X-100 in PBS with 4% normal goat serum overnight at room temperature. Slices were washed the next day, 4 × 10 min each in PBS, mounted with a drop of Prolong Gold mounting media (Thermo Fisher Scientific) on Superfrost Plus slides, using standard coverslips. Slides were allowed to air dry in the dark at room temperature before being imaged for morphological analysis. DnKv4.2 transfected neurons were confirmed using the red channel on the Olympus FV1000 laser-scanning confocal microscope. Filled neurons green (dnKv4.2-negative) and green/red (dnKv4.2-positive) were imaged at multiple Z-planes using a two-photon microscope (Olympus FV1000-MPE) with a multiimmersion objective lens (ULTRA 25x, NA 1.05, Resolution: 3.1447 pixels per micron, Voxel size: 0.3180 x 0.3180 x 0.98 micron^^3^). Imaged neurons were reconstructed using semiautomatic settings in Neuromantic software. Apical and basal length and nodes were automatically computed by the software and compared across the genotypes.

### Statistical Analysis

Data analyses were conducted in GraphPad Prism 8 (GraphPad Software). To determine the distribution and the homogeneity of variance, all datasets were explored by using D’Agostino & Pearson test and Kolmogorov-Smirnov test, respectively. All data are expressed as mean ± SEM and statistical significance was determined at *p*-values <0.05. For genotype comparisons of L2/3 neurons, two-sided Student’s t-test were used for normally distributed data and Mann Whitney U tests when the data did not follow a normal distribution. In some electrophysiology analyses, two-way RM-ANOVA with Šídák’s correction for post-hoc multiple comparisons were used to compare the number of spikes in response to current injections, while a standard two-way ANOVA with Tukey’s multiple comparisons post-hoc test was used to compare measurements between two groups (genotypes) when treatment was included in the experiment. See figure legends for specific details on statistical tests for each experiment.

## Figure Legends

**Supplemental Figure 1:**
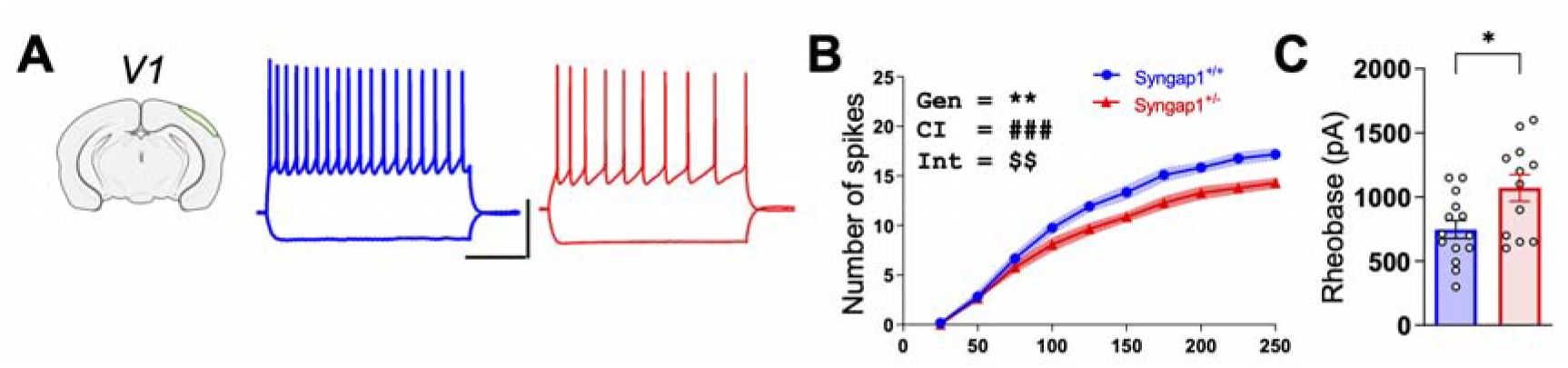
*Syngap1* regulates IME in developing V1 neurons. **(A)** Representative current clamp traces from L2/3 V1 excitatory neurons from acute Syngap1^+/+^ and Syngap1^+/-^ coronal slices, Scale bar 40mV, 200 ms). **(B)** The I/O plot shows significantly reduced AP firing with increasing current injection [Repeated measures ANOVA, Genotype, F (1, 24) = 13.31, p=0.0013; Injected current x Genotype Interaction F (10, 240) = 2.954, p<0.0016; Injected current F (10.240) = 330.8, p<0.0001. *Syngap1*^+/+^ = 3 mice, 14 neurons; *Syngap1*^+/-^ = 3 mice, 12 neurons]. **(C)** Scatter plots shows significantly increased rheobase in the same set of L2/3 *Syngap1*^+/-^ V1 neurons as in (B) (unpaired t test, t(24)=2.649, p=0.014). Bar graphs represent mean ± SEM, *p < 0.05, **p<0.01, ***p<0.001. For line graphs, symbols mean neuron means, area fill indicate SEMs, and in RM-ANOVA, main effects are denoted by the following symbols: * represents genotype, # represents current injection and $ represents genotypic x injected current interaction.

**Supplemental Figure 2:**
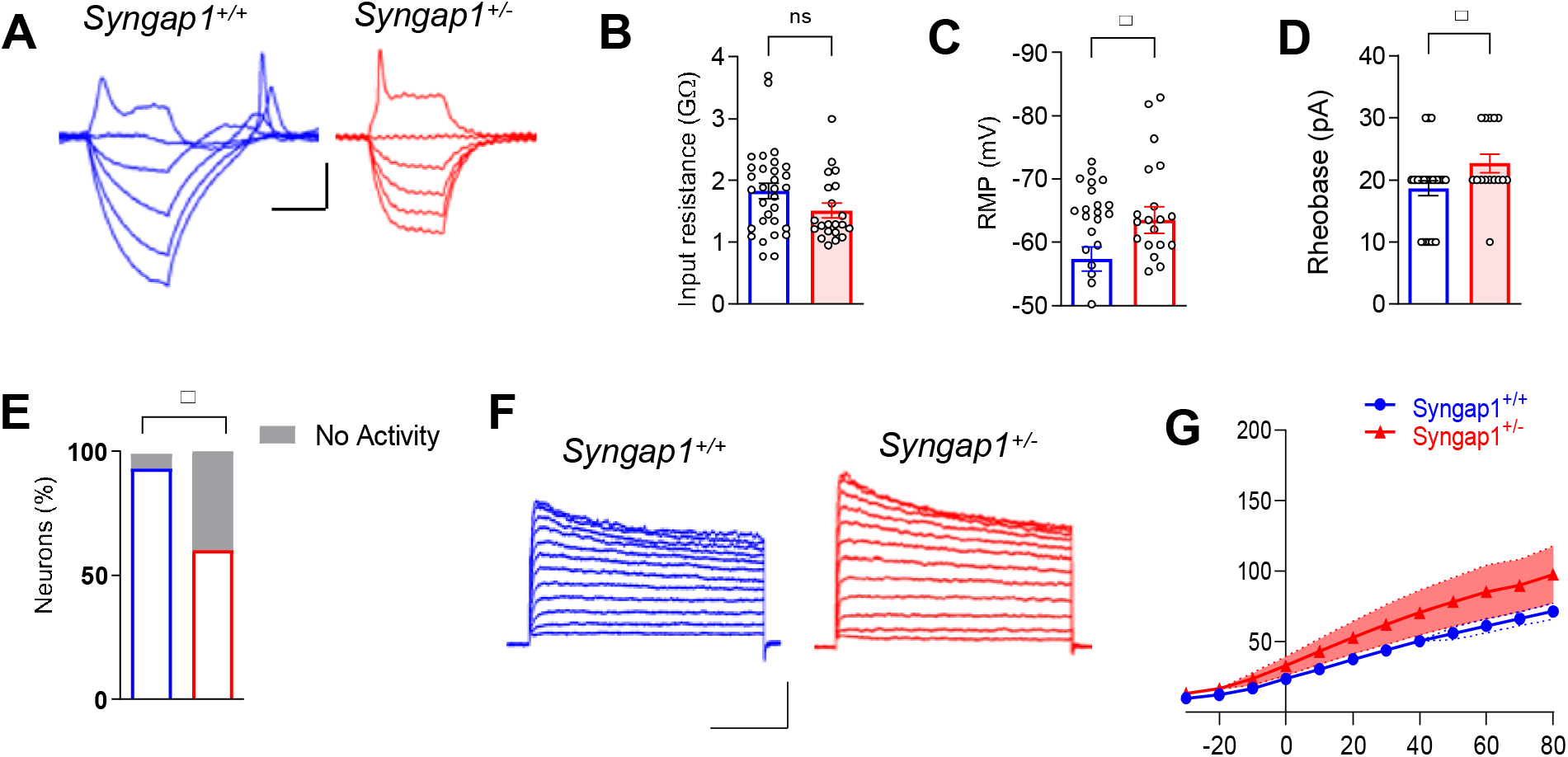
*Syngap1* regulates pyramidal cell excitability in newborn animals. (**A**) Representative traces for membrane potential changes to 150 ms current pulses from −100 pA to+40 pA in steps of 10 pA to test firing ability, resting membrane potential, rheobase and input resistance of postnatal cortical plate (CP) neurons at PND 0-1 from *Syngap1*^+/+^ (blue) or *Syngap1*^+/-^ (red) acute slices (Scale bar 20mV, 100 ms). (**B**) Resting membrane potential was measured for CP neurons (unpaired t-test, t(50)=2.129 p=0.0380; *Syngap1*^+/+^ =31 neurons from 5 mice, *Syngap1*^+/-^ =21 neurons from 3 mice). (**C**) Input resistance from same neuronal populations as in (B); t (50) =1.740, p=0.0881; unpaired t test; *Syngap1*^+/+^ =31 neurons from 5 mice, *Syngap1*^+/-^ =21 neurons from 3 mice. (**D**) Rheobase determined as first step required to elicit an action potential spike from the same neuronal population as in (B); t (42) =2.037, p=0.0480; unpaired t test; *Syngap1*^+/+^ =29 neurons from 5 mice, *Syngap1*^+/-^ =15 neurons from 3 mice. (**E**) Percentage of successful observations of spiking activity in CP neurons from *Syngap1*^+/+^ (blue) or *Syngap1*^+/-^ (red) acute slices (p<0.0001; Fischer’s exact test; *Syngap1*^+/+^ =31 neurons from 5 mice, *Syngap1*^+/-^ =21 neurons from 3 mice). (**F**) Representative potassium channel current waveforms in response to 300ms depolarizing pulse of varying test potentials from holding potential of −30mV to +70mV from postnatal CP *Syngap1*^+/+^ or *Syngap1*^+/-^ neurons (Scale bar 500 pA, 100ms). (**G**) Peak potassium current densities for L2/3 neurons are plotted as a function of test potential; RM 2-way ANOVA, Genotype F (1.19) =2.162, p<0.1578; Test potential x Genotype Interaction F (11,209)=1.558, p<0.1132; Test potential F (11.209)=61.90, p<0.0001; n=12 neurons from 3 *Syngap1*^+/+^ mice and n=9 neurons from 3 *Syngap1*^+/-^ mice.

**Supplementary Figure 3:**
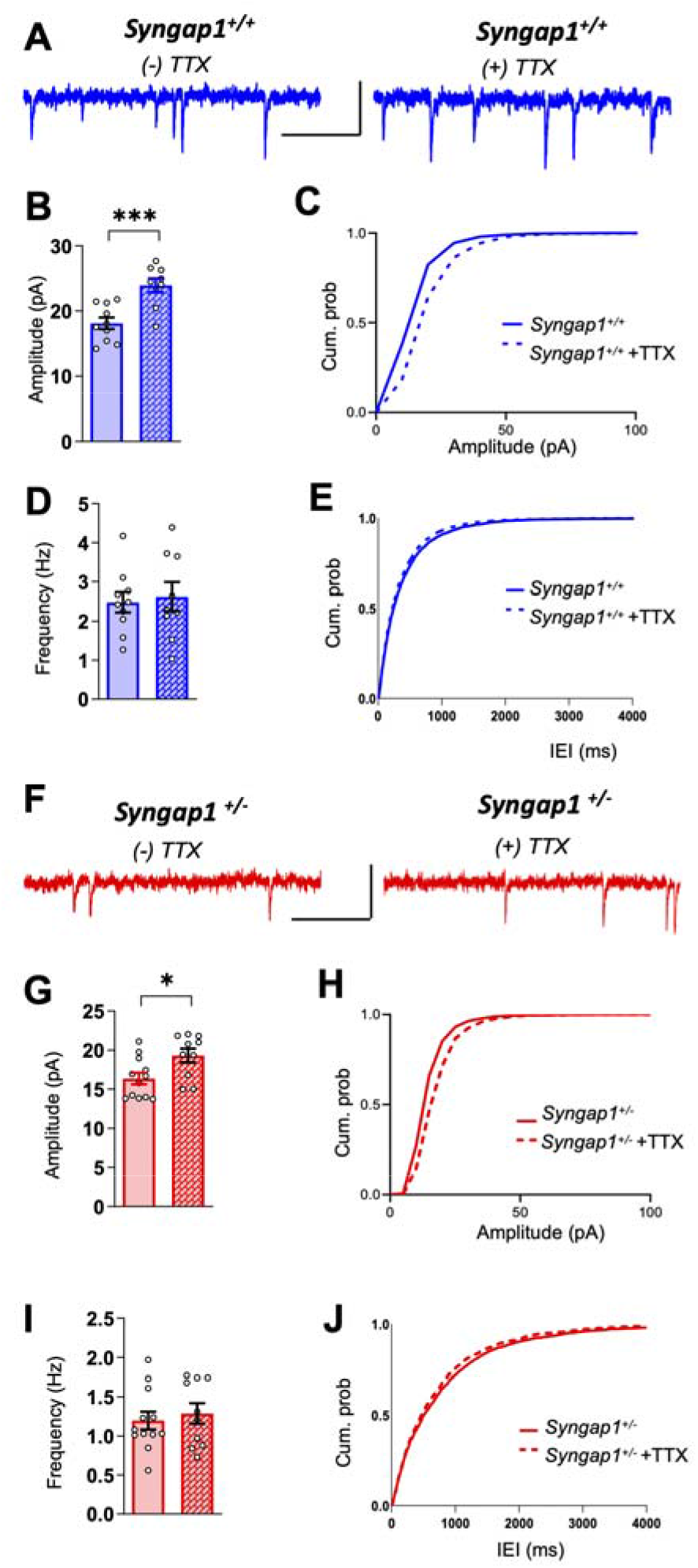
Homeostatic plasticity of excitatory synapse strength persists in SSC layer 2/3 excitatory neurons from *Syngap1*^+/-^ mice. **(A)** Representative mEPSC responses from L2/3 SSC cells in organotypic cultures from untreated and TTX (1 μM) treated *Syngap1*^+/+^ organotypic cultures (Scale bar 40 pA, 1s). **(B)** Quantification of mEPSC amplitudes for L2/3 SSC cells in *Syngap1*^+/+^ organotypic cultures subjected to each condition in (A) (unpaired t test, t(17) = 4.233, p=0.0006; n=10 cells from 4 slices (2 mice) for *Syngap1*^+/+^, n=9 cells from 3 slices (2 mice) for *Syngap1*^+/+^ + TTX). **(C)** Cumulative probability histograms of mEPSC amplitudes in *Syngap1*^+/+^ neurons from untreated and TTX treated organotypic cultures. (**D**) Quantification of frequency of mEPSCs for L2/3 SSC cells in organotypic cultures subjected to each condition in (A) (unpaired t test, t(17)=0.3278, p=0.7470; n=10 cells from 4 slices (2 mice) for *Syngap1*^+/+^ n=9 cells from 3 slices (2 mice) for *Syngap1*^+/+^ + TTX). **(E)** Cumulative probability histograms of mEPSCs interevent interval for *Syngap1*^+/+^ neurons from untreated and TTX treated organotypic cultures. (**F**) Representative mEPSC responses from L2/3 SSC cells from untreated and TTX (1 μM) treated *Syngap1*^+/-^ organotypic cultures (Scale bar 40 pA, 1s). (**G**) Quantification of mEPSC amplitudes for L2/3 SSC cells in *Syngap1*^+/-^ organotypic cultures subjected to each condition in (F) (unpaired t test, t(20) = 2.594, p=0.0238; n=12 cells from 4 slices(3 mice) for *Syngap1*^+/-^, n=10 cells from 4 slices (4 mice) for *Syngap1*^+/-^ + TTX). (**H**) Cumulative probability histograms of mEPSC amplitudes for *Syngap1*^+/-^ neurons from untreated and TTX treated organotypic cultures. (**I**) Quantification of frequency of mEPSCs for L2/3 SSC cells in organotypic cultures subjected to each condition in (F) (unpaired t test, t(20)=0.5195, p=0.6091; n=12 cells from 4 slices (3 mice) for *Syngap*^+/-^, n=10 cells from 4 slices (4 mice) for *Syngap1*^+/-^ + TTX). (**J**) Cumulative probability histograms of mEPSC interevent interval for *Syngap1*^+/-^ neurons from untreated and TTX treated organotypic cultures. Bar graphs represent mean ± SEM, *p < 0.05, **p<0.01, ***p<0.001.

**Supplemental Figure 4:**
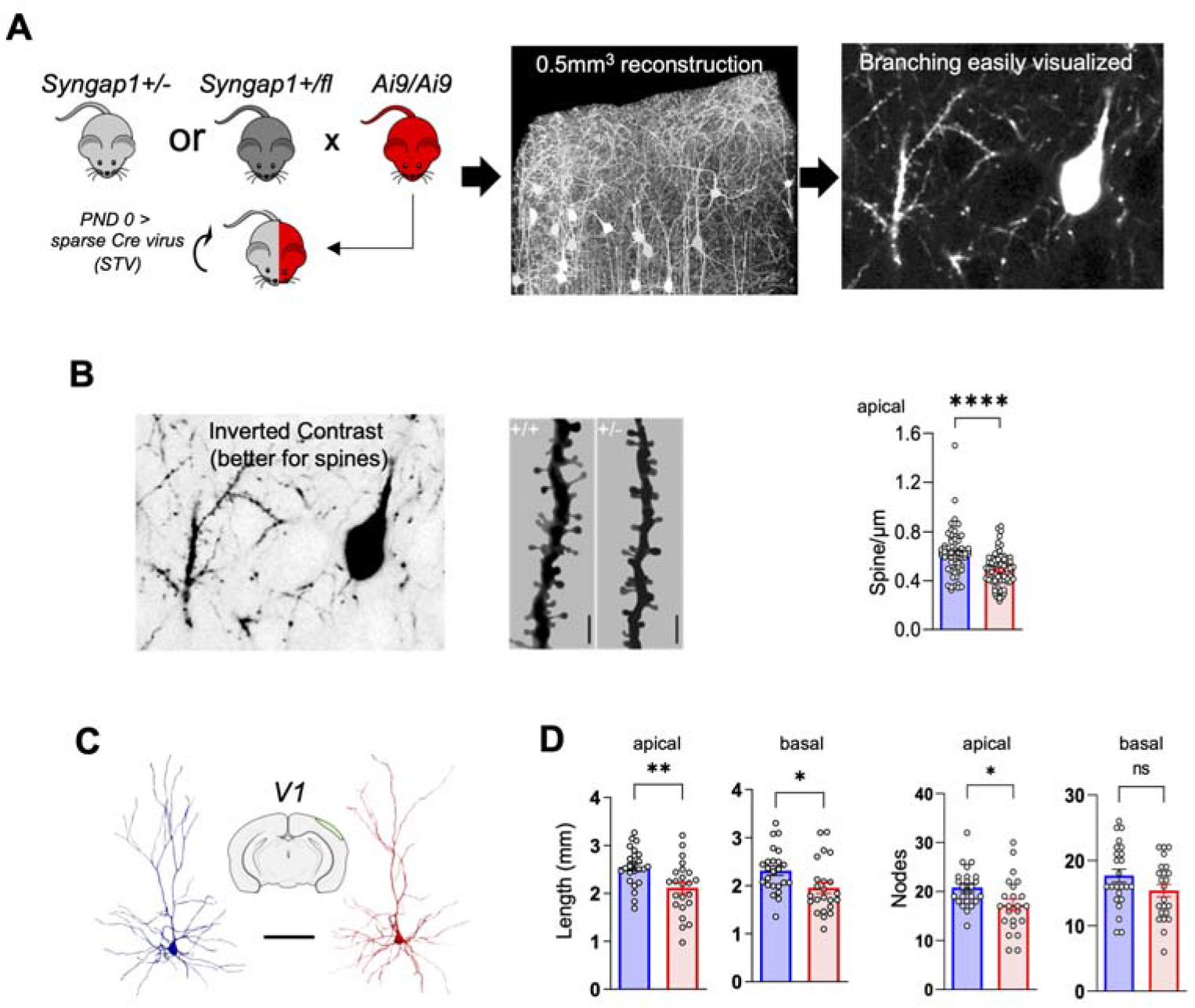
Dendritic morphogenesis methodology and morphology of V1 L2/3 neurons. **(A)** Experimental design showing crosses of *Syngap1* strains with TdTomato Ai9 Cre reporter line (#007905, Jackson Lab) followed by Superior temporal vein (STV) injections of AAV9-Cre to induce TdTomato expression in a sparse population of neurons in *Syngap1* cKO mice. **(B)** Spine density was measured in apical dendrites from L2/3 of SSC neurons of P14 animals. Image on the left shows inverted contrast image for better visualization of dendritic spines and on the right is representative apical branches from layer 2/3 neurons from *Syngap1*^+/+^ and *Syngap1*^+/-^ mice with quantification of apical dendritic spine density (spine density – Mann-Whitney’s exact test: U=1283, p< 0.0001; *Syngap1*^+/+^= 68 dendritic segments from 4 mice, *Syngap1*^+/-^ = 77 dendritic segments from 5 mice). **(C)** Representative 3D reconstructions of apical and basal branches of L2/3 neurons from primary visual cortex (V1), in *Syngap1*^+/+^ (Blue) and *Syngap1*^+/-^ (Red) mice (Scale bar 100 μm). **(D)** Bar histograms showing morphological parameters of V1 apical (total length - unpaired t-Test, t(45)=3.102, p = 0.0033; # of nodes, t (45)=2.493, p=0.0164; *Syngap1*^+/+^ = 4 mice, 24 neurons; *Syngap1*^+/-^ = 4 mice, 23 neurons) and basal (total length – unpaired t-Test, t(45)=2.402, p = 0.0205; # of nodes, t(45)=1.665, p = 0.1028; *Syngap1*^+/+^ = 4 mice, 23 neurons; *Syngap1*^+/-^ = 4 mice, 24 neurons) dendritic arbors. Values represent means; error bars indicate SEMs.

**Supplemental Figure 5:**
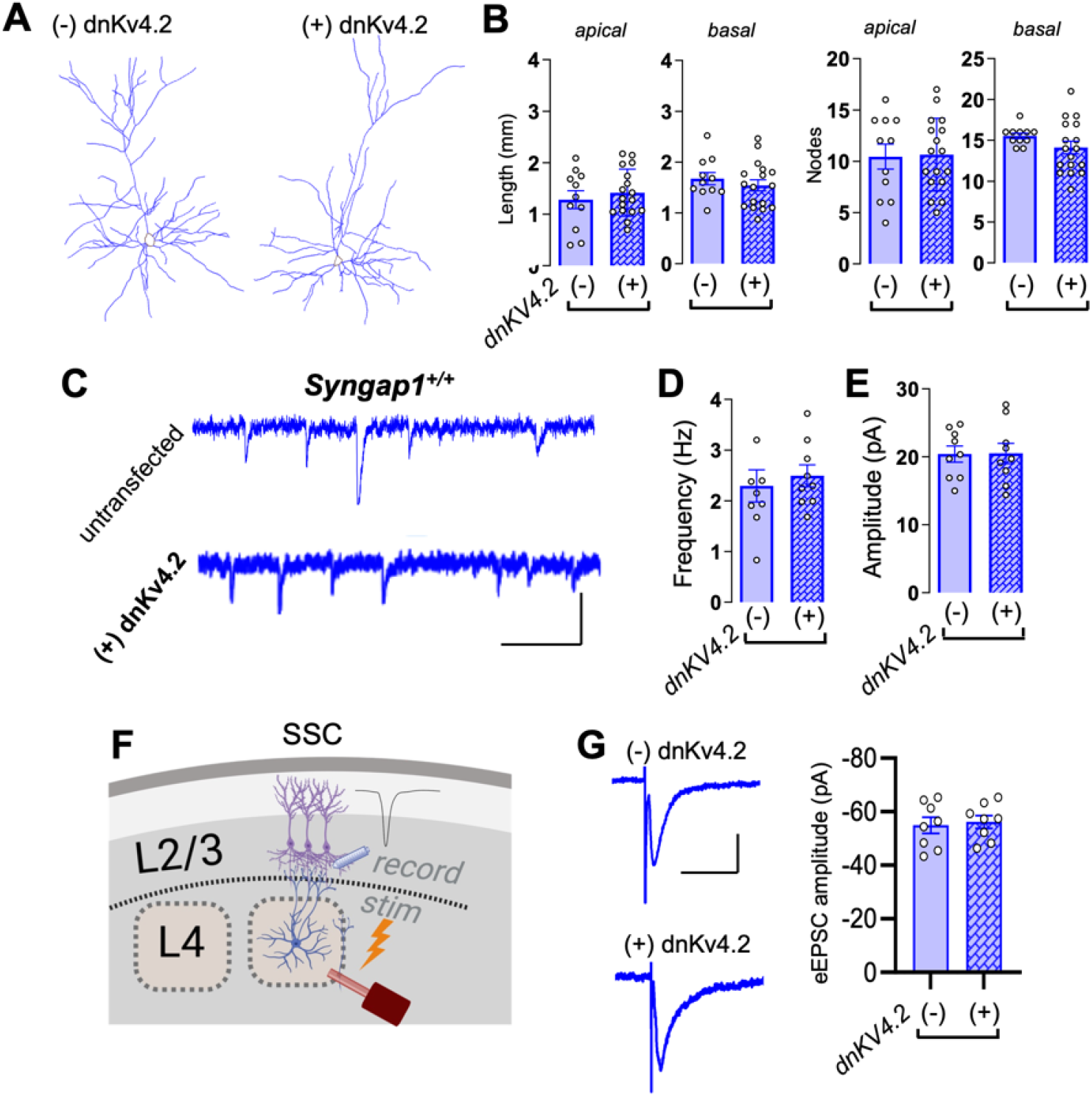
Enhancing IME in *Syngap1*^+/+^ SSC neurons has no effect on dendritic morphogenesis or excitatory circuit connectivity. **(A)** Representative reconstructions of neurobiotin filled, Streptavidin-Alexa Fluor 488 labelled (-) dnKv4.2 untransfected and (+) dnKv4.2 transfected L2/3 SSC neurons from acute coronal slices prepared from *Syngap1*^+/+^ mice; scale bar 100 μm. (**B**) Quantification of apical and basal dendritic length and nodes of traced (-) dnKv4.2 and (+) dnKv4.2 neurons from *Syngap1*^+/+^ mice (Apical dendrites, total length – unpaired t test t(26)=0.6914, p=0.4954; # of nodes-unpaired t test t(26)=0.1329, p=0.8953; Basal dendrites, total length – Unpaired t test t(26)=0.7821, p=0.4412; # of nodes unpaired t test, t(26)=1.408, p=0.1709; *Syngap1*^+/+^ = 2 mice, (-)dnKv4.2=11 neurons, (+)dnKv4.2=17 neurons]. (**C**) Representative mEPSCs traces from (-) dnKv4.2 untransfected and (+) dnKv4.2 transfected L2/3 SSC neurons from PND12-14 *Syngap1*^+/+^ acute slices; Scale bar 1s, 40pA. (**D**) Bar histograms showing quantification of mEPSCs frequencies from (-) dnKv4.2 untransfected and (+) dnKv4.2 transfected L2/3 SSC neurons from PND12-14 *Syngap1*^+/+^ mice (unpaired t test t (16) =0.5215; p=0.6092). **(E)** mEPSC amplitudes from same neurons (unpaired t test t (16) =0.05157, p=0.9595). **(F)** Cartoon depicting experimental setup for measuring feed-forward excitation from L4>L2/3 excitatory neurons. **(G)** *Left* - Representative L4-evoked L2/3 EPSC traces from untransfected or transfected L2/3 excitatory neurons from acute *Syngap1*^+/+^ slices, Scale bar 20 pA, 50 ms. **(G)** *Right* - Bar histogram plot of evoked EPSC amplitudes in these populations (unpaired t test, t(14)=0.3428, p=0.7369) Bar graphs represent mean ± SEM, *p < 0.05, **p<0.01, ***p<0.001.

**Figure S6:**
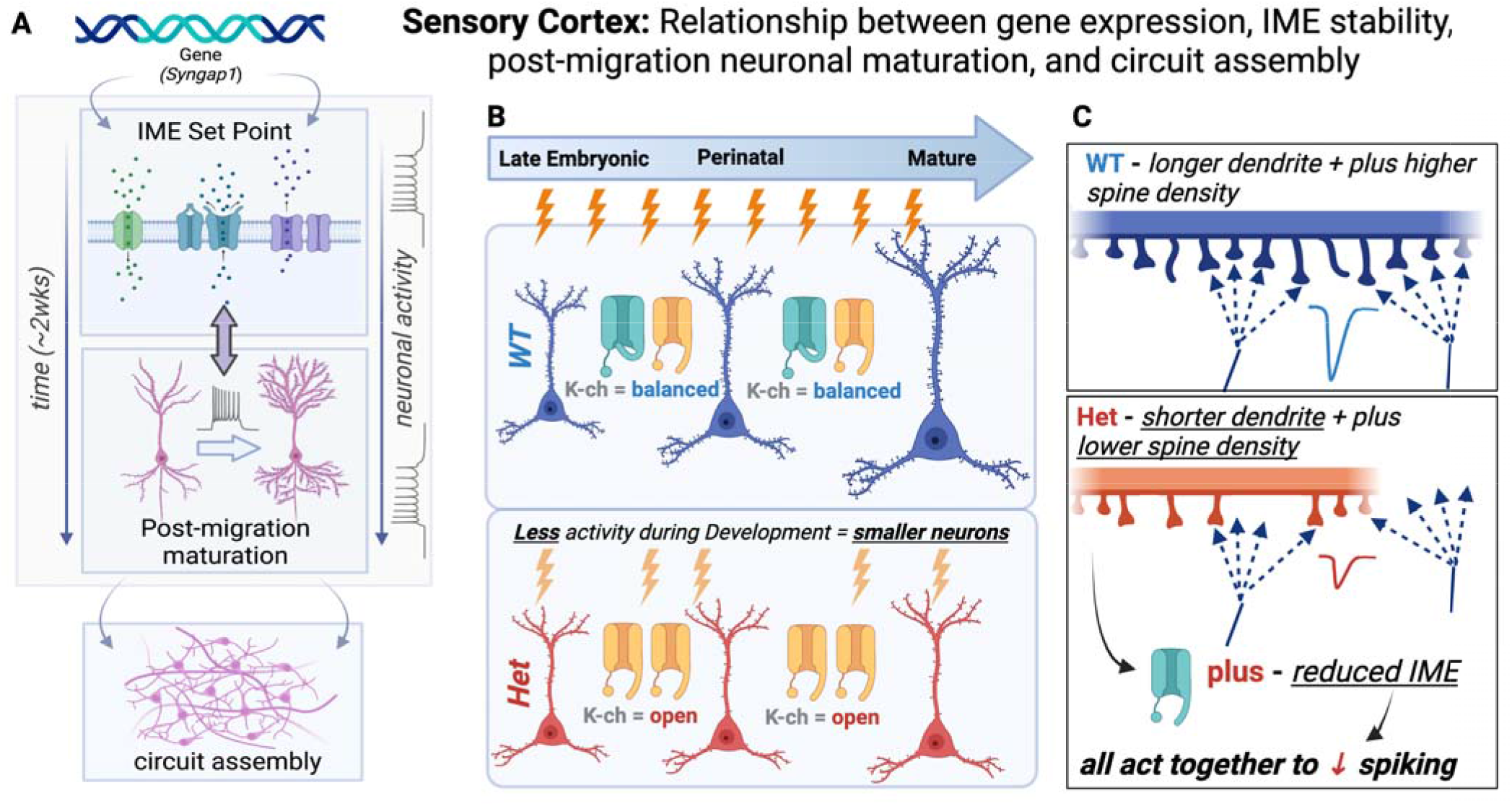
Experimental model based on outcome of the current studies. **(A)** Cartoon depicting relationship of *Syngap1* expression, the importance of IME stability in L2/3 during the early postnatal period, dendritic maturation, and assembly of cortical circuits. **(B-C)** Cartoons depicting the role of *Syngap1* on regulation of ion channels that are known to control IME and how regulation of these ion channels contribute to dendritic maturation and synaptic connectivity that underlies cortical circuit assembly.

## Notes

### Competing Interest Statement

The authors have declared no competing interest.

